# Dual-targeted transcription factors are required for optimal photosynthesis and stress responses in *Arabidopsis thaliana*

**DOI:** 10.1101/793968

**Authors:** Piotr Gawroński, Paweł Burdiak, Lars B. Scharff, Jakub Mielecki, Magdalena Zaborowska, Cezary Waszczak, Stanisław Karpiński

**Affiliations:** Department of Plant Genetics, Breeding, and Biotechnology, Warsaw University of Life Sciences, 02–776 Warsaw, Poland; Copenhagen Plant Science Center, Department of Plant and Environmental Sciences, University of Copenhagen, 1871 Frederiksberg C, Denmark; Organismal and Evolutionary Biology Research Programme, Faculty of Biological and Environmental Sciences, and Viikki Plant Science Centre, University of Helsinki, 00014 Helsinki, Finland

**Keywords:** Chloroplast retrograde signaling, CIA2, CIL, nonphotochemical quenching, photosynthesis, thermo- and photooxidative stress tolerance, chloroplast translation

## Abstract

Chloroplast to nucleus retrograde signaling is essential for cell function, acclimation to fluctuating environmental conditions, plant growth and development. The vast majority of chloroplast proteins are nuclear-encoded and must be imported into the organelle after synthesis in the cytoplasm. This import is essential for the development of fully functional chloroplasts. On the other hand, functional chloroplasts act as sensors of environmental changes and can trigger acclimatory responses that influence nuclear gene expression. Signaling *via* mobile transcription factors (TFs) has been recently recognized as a way of communication between organelles and the nucleus. In this study, we performed a targeted reverse genetic screen to identify novel dual-localized TFs involved in chloroplast retrograde signaling during stress responses. We found that CHLOROPLAST IMPORT APPARATUS 2 (CIA2), a TF with putative plastid transit peptide can be detected in chloroplasts and the nucleus. Further, we found that CIA2, along with its homolog CIA2-like (CIL) act in an unequally redundant manner and are involved in the regulation of Arabidopsis responses to UV-AB, high light, and heat shock. Finally, our results suggest that both CIA2 and CIL are crucial for chloroplast translation. Our results contribute to a deeper understanding of signaling events in the chloroplast-nucleus cross-talk.

**Significance:** We found that a transcription factor CIA2 can be located in chloroplasts and nucleus. CIA2 and is close homolog CIL are involved in protein translation and abiotic stress responses, and we suggest that they play an essential role in retrograde signaling between these organelles.

## Introduction

In plants, intracellular communication between the nucleus, chloroplasts, and mitochondria is essential for the regulation and coordination of physiological processes such as growth, development, stress responses, photosynthesis, and respiration (de Souza *et al.*, 2017). Mechanisms that coordinate organellar and nuclear gene expression enable responses to fluctuating or rapidly changing environmental conditions.

Chloroplasts contain a few thousand proteins that are involved in photosynthesis, intracellular signaling, and biosynthesis of fatty acids, amino acids, hormones, vitamins, nucleotides, and secondary metabolites (Leister and Kleine, 2008). Most of these proteins are encoded by the nuclear genome, synthesized by cytosolic ribosomes, imported into the chloroplast, and targeted to a specific compartment within the chloroplast (Jarvis, 2008). Chloroplast genomes encode about 80 of these proteins, most of which function in photosystems, photosynthetic electron transport, and the organellar gene expression machinery (Scharff and Bock, 2014; Daniell *et al.*, 2016). Consequently, dark and light reactions of photosynthesis—the function of the chloroplast—are strictly dependent on the interorganelle communication as the components of the photosynthetic apparatus are encoded by nuclear and plastid genomes, and a close cooperation between these two genomes is required for the development of functional chloroplasts and chloroplast plasticity (Jarvis and López-Juez, 2013).

The functionality of the photosynthetic apparatus depends on factors such as water and nutrient supply, light, and temperature (Gururani *et al.*, 2015). Besides photosynthesis, chloroplasts play an important role as redox sensors of environmental conditions and trigger acclimatory responses (Z., Li *et al.*, 2009). Changes in the developmental and metabolic states of chloroplasts or in the redox status of photosynthetic electron carriers can trigger alterations in the nuclear gene expression in a process called retrograde signaling (Chi *et al.*, 2013; Estavillo *et al.*, 2013; Guo *et al.*, 2016). It is now well established that the perturbation of multiple plastid processes, including tetrapyrrole biosynthesis, protein synthesis, reactive oxygen species (ROS) metabolism, and dark and light reactions of photosynthesis, influences the expression of nuclear genes encoding photosynthetic proteins (Pesaresi *et al.*, 2006). Moreover, chloroplast retrograde signaling not only coordinates the expression of nuclear and chloroplast genes, which is essential for chloroplast biogenesis, but also ensures chloroplast vitality in changing environmental conditions (Barajas-López *et al.*, 2013) and triggers the expression of nuclear-encoded genes for other cellular compartments such as the cytoplasm and peroxisomes (Karpinski *et al.*, 1997; Karpiński *et al.*, 1999; Mateo *et al.*, 2004; Mühlenbock *et al.*, 2008; Pogson *et al.*, 2008).

Changes in the absorbed light quality and intensity result in rapid changes in the redox state of photosynthetic electron carriers and lead to unbalanced production of ROS such as hydrogen peroxide (H_2_O_2_), singlet oxygen, and superoxide anions (Karpiński *et al.*, 1999; Mullineaux *et al.*, 2006; Mühlenbock *et al.*, 2008; Pogson *et al.*, 2008). Recently, it was proposed that H_2_O_2_ produced in chloroplasts can be directly transported to the nucleus to act as a signaling molecule as its accumulation in both compartments was observed immediately after exposure to light (Caplan *et al.*, 2015; Exposito-Rodriguez *et al.*, 2017). Moreover, H_2_O_2_ produced in chloroplasts under drought and excessive light conditions influences the metabolism of 3’-phosphoadenosine 5’-phosphate (PAP), which, upon accumulation, modulates the expression of nuclear stress-responsive genes (Estavillo *et al.*, 2011; Chan *et al.*, 2016). Increased singlet oxygen generation in chloroplasts can also trigger specific retrograde signals. However, due to the high reactivity of singlet oxygen, its half-life is too short to enable direct transport to the nucleus, and it was proposed that carotenoid oxidation product, β-cyclocitral, acts as a stress signal induced by singlet oxygen produced in grana stacks (Ramel *et al.*, 2012). Moreover, singlet oxygen can be also produced in grana margins where it induces retrograde signaling through two plastid localized proteins, EXECUTER 1 and 2 (Lee *et al.*, 2007; Wang *et al.*, 2016; Dogra *et al.*, 2019).

In addition to signaling via ROS and metabolites, many transcription factors (TFs) were shown to be controlled by signals generated in the organelles. There are two known TFs, ANAC013 and ANAC017, which respond to the mitochondrial redox status (De Clercq *et al.*, 2013; Ng *et al.*, 2013). These TFs are anchored in the endoplasmic reticulum membrane, and in response to signals from mitochondrial complex III, they are released to the nucleus by the proteolytic cleavage of their transmembrane domains. After translocation to the nucleus, ANAC013 and ANAC017 regulate the expression of mitochondrial dysfunction stimulon genes (De Clercq *et al.*, 2013; Ng *et al.*, 2013). Further, it was recently shown that RADICAL-INDUCED CELL DEATH1 (RCD1) interacts with ANAC013 and ANAC017 to integrate ROS signals from chloroplasts and mitochondria (Shapiguzov *et al.*, 2019). Thus, the dual localization of TFs presents a possibility of their function in retrograde signaling.

Early, *in silico* analyses of Arabidopsis genes encoding putative TFs predicted targeting of at least 48 TFs to the plastids (Wagner and Pfannschmidt, 2006). Later, another *in silico*-based screen approach predicted that 78 Arabidopsis TFs reside in the plastids (Schwacke *et al.*, 2007). Indeed, several proteins exhibiting dual nuclear–plastid localization might potentially be involved in signal transduction pathways involving regulatory protein storage in the plastids. It was shown that most of the dual-targeted (nucleus and organelle) proteins have functions in the maintenance of DNA, telomere structuring, gene expression, or innate immunity (Krause *et al.*, 2012; Caplan *et al.*, 2015). These *in silico* studies were supported by *in vivo* evidence with WHIRLY1 being the first protein to be identified in the nucleus and plastids of the same plant cell (Grabowski *et al.*, 2008); however, its molecular function remains elusive. Later, the plant homeodomain (PHD) transcription factor PTM (PHD-type TF with transmembrane domains) was shown to accumulate in the nucleus after release from the plastid surface. In the nucleus, PTM is thought to activate the transcription factor ABA INSENSITIVE 4 (ABI4), thereby providing a way to communicate the plastid status to the nucleus (Sun *et al.*, 2011). However, the roles of PTM and ABI4 in chloroplast-to-nucleus communication were recently questioned (Page *et al.*, 2017; Kacprzak *et al.*, 2019).

Although nuclear gene expression involves transcriptional control, it is generally believed that in the course of evolution, the regulation of plastid gene expression has shifted from predominantly transcriptional to predominantly posttranscriptional (Eberhard *et al.*, 2002), although significant transcriptional regulation occurs in chloroplasts (Liere and Börner, 2007). Posttranscriptional control is exerted at the level of mRNA stability and, most importantly, at the level of mRNA translation (Zoschke and Bock, 2018). During chloroplast biogenesis, translational regulation is required for the differentiation of chloroplasts from proplastids during the early development of stem and leaf tissues (Sugiura, 2014). At the later stages, in mature chloroplasts, translation is regulated mostly by light and controls chloroplast growth for division in the expanding cells of green tissues. Translational regulation also occurs in response to changing environmental conditions and quite often is essential to repair the photosynthesis machinery (Chotewutmontri and Barkan, 2018). For example, the regulation of *psbA* translation encoding for D1 core protein of photosystem II (PSII) is crucial for the repair of photodamaged PSII complexes, whereas the repression of *rbcL* translation encoding for a large subunit of RuBisCo occurs during oxidative stress (Nickelsen *et al.*, 2014).

In this paper, we aimed to characterize dual-localized TFs that can be involved in the communication between the chloroplast and the nucleus. As a result of a targeted reverse genetic screen, we focused on CHLOROPLAST IMPORT APPARATUS 2 (CIA2) and its homolog CIA2-like (CIL), which encode TFs with conserved *CONSTANS*, *CO-like*, and *TOC1* (CCT) domain at the C-terminus and putative plastid transit peptide at the N-terminus. Our results suggest that CIA2 and CIL play an important role in the regulation of chloroplast translation, thus influencing photosynthetic electron transport, accumulation of photosynthetic pigments, and chloroplast retrograde stress responses in *Arabidopsis thaliana*.

## Results

### Reverse genetic screen to identify chloroplast-targeted TFs involved in retrograde signaling

Because chloroplast-to-nucleus signaling pathways are not sufficiently understood, we decided to use the targeted reverse genetic screen to identify new retrograde signaling components. To this end, we focused on the identification of dual-localized TFs due to their potential role in chloroplast retrograde signaling (Sun *et al.*, 2011). To identify such TFs, we extracted and compared the gene identifiers obtained from three *Arabidopsis thaliana* plastid proteome databases (Schwacke *et al.*, 2003; Q., Sun *et al.*, 2009; Myouga *et al.*, 2013) and two transcription factor databases (Pérez-Rodríguez *et al.*, 2009; Jin *et al.*, 2014). This allowed us to identify a set of TFs with potential chloroplast localization. Further, we obtained T-DNA insertion lines for a selected set of genes (Data S1). These mutants were then subjected to conditions promoting chloroplast photooxidative stress, as we hypothesized that mutants lacking efficient communication between the chloroplast and the nucleus (possibly dependent on TFs) would show altered susceptibility to such treatments. During the primary screen, two types of stress treatments were applied: illumination with ultraviolet AB (UV-AB) and exposure to high light intensity in combination with cold (cHL). UV-AB significantly impairs photosynthetic electron transport and the general function of photosynthetic machinery (Hollósy, 2002; Caldwell, 1993). At the molecular level, UV-AB damages the ribosomes by crosslinking the cytosolic and chloroplast ribosomal proteins to RNA, thereby transiently inhibiting translation *in vivo* (Casati and Walbot, 2004; Ferreyra *et al.*, 2010). On the other hand, cHL causes imbalances in photosynthetic reactions resulting in the photoinhibition of photosystem II (PSII) and oxidative stress (Yabuta *et al.*, 2002; Distelbarth *et al.*, 2013). Most of the analyzed mutants were not affected by treatment with UV-AB and cHL as compared to Col-0 (Figure S1); however, based on the phenotypes of mutant lines, we could select eight candidate genes for further investigation (Table S1). Next, the coding sequences of six of the eight candidate genes were cloned, and fusion proteins with C-terminal YFP were expressed under the control of 35S promoter in Arabidopsis cotyledons to confirm their putative chloroplast localization (Figure S2 and Table S1). Except for p35S::AT5G57180:YFP, which showed chloroplast and nuclear localization, all the analyzed fusion proteins were found exclusively in the nucleus (Figure S2, Table S1). AT5G57180 encodes CHLOROPLAST IMPORT APPARATUS2 (CIA2) protein, which was previously found to play a role in protein import into chloroplasts (Sun *et al.*, 2001) and protein synthesis in chloroplasts (C.,-W., Sun *et al.*, 2009). In our reverse genetic screening (Figure 1a,b), line SALK_045340 (hereafter referred to as *cia2–4*) that harbors T-DNA insertion in the first intron of the CIA2 gene (Figure 1c) showed increased susceptibility to UV-AB as indicated by the lower maximum efficiency of PSII (*F*_v_/*F*_m_) and higher ion leakage resulting from the induction of cell death (Figure 1a). Similarly, the exposure to cHL reduced the *F*_v_/*F*_m_ of *cia2–4* to a higher extent than that in the exposure to Col-0 (Figure 1b). The CIA2 protein contains a C-terminal conserved CCT motif, a characteristic feature of one family of TFs involved in light signal transduction (Strayer *et al.*, 2000). The CCT domain contains a putative nuclear localization signal (NLS), which is consistent with the nuclear localization of CIA2 observed earlier (Sun *et al.*, 2001). However, our *in silico* analysis of CIA2 protein sequence predicted the existence of an N-terminal 59 amino acid-long chloroplast transit peptide, which suggests that CIA2 can be imported into chloroplasts (Figure 1c and Data S1) (Schwacke *et al.*, 2003; Myouga *et al.*, 2013). The expression of full-length CIA2 fused with YFP (p35S::CIA2:YFP) resulted in a weak fluorescence signal; thus, we also expressed a 100 amino acid-long N-terminal part of CIA2 fused N-terminally to YFP (p35S::CIA2^1–100^:YFP). As a result, we observed a notably higher fluorescence intensity and confirmed the function of CIA2 transit peptide as the presence of fusion protein was detected in chloroplasts (Figure S2). Thus, based on the initial screen and protein localization experiments, we decided to focus on CIA2 that appears to be localized in the nucleus and chloroplast and is required for acclimatory responses to UV-AB and cHL.

**Figure 1.**
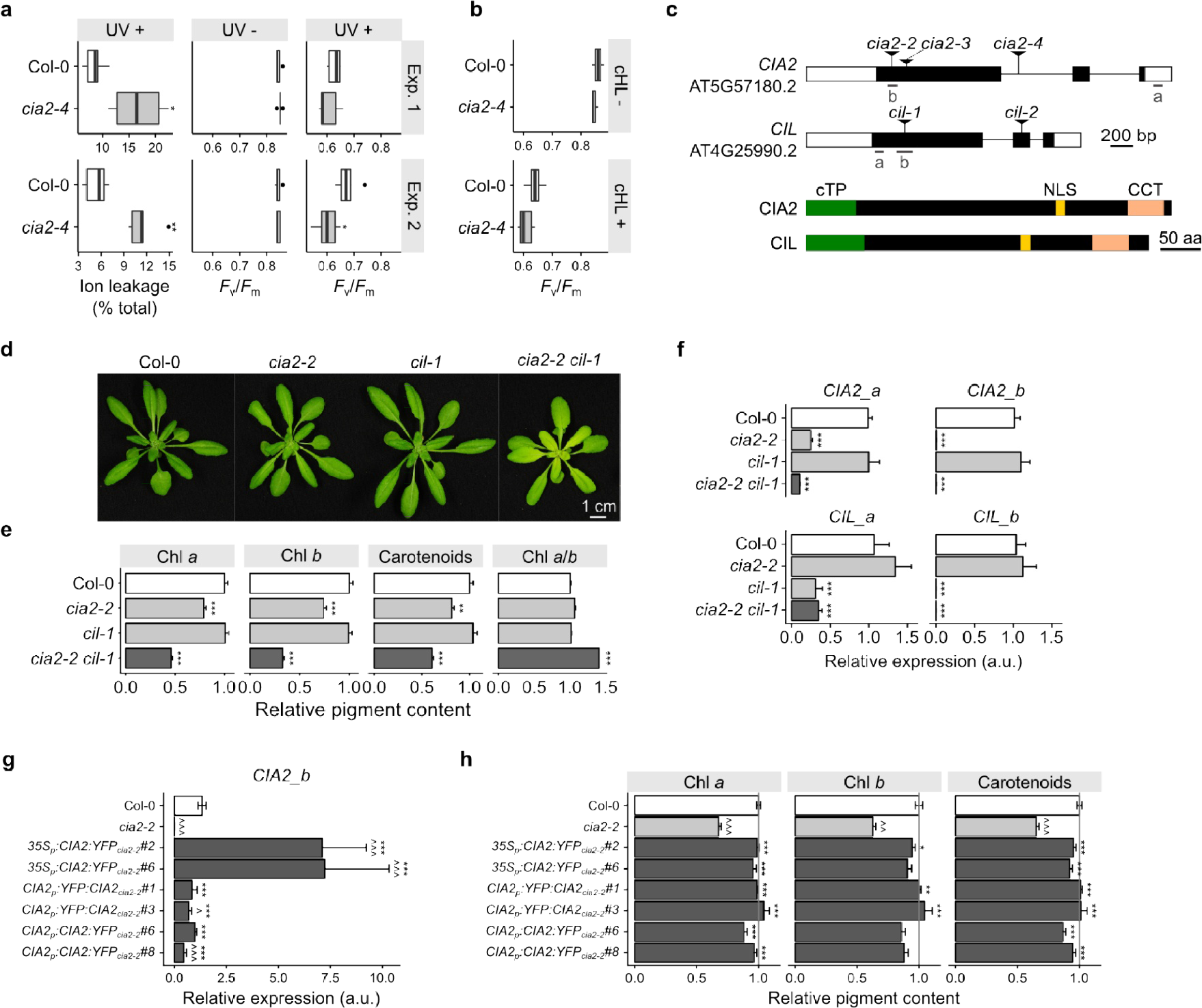
Isolation and characterization of *cia2–2 cil-1* double mutant. (**a**, **b**) Isolation of *cia2–4* mutant in reverse genetic screening using UV-AB (**a**) and high light in combination with low temperature (cHL) (**b**). (**c**) In the upper part, a schematic representation of *CIA2* and *CIL* genes is presented. Black and white rectangles represent exons and untranslated regions, respectively. The lower part shows the analysis of functional domains in CIA2 and CIL proteins: cTP–chloroplast transit peptide, NLS–nuclear localization signal, CCT–putative active domain in CIA2 and CIL. The CCT domain contains NLS; however, it is not shown in the diagram for simplicity. (**d**) Phenotypes of 4-week-old plants grown in long-day conditions. *cia2–2* and *cia2–2 cil-1* plants were paler than the rest of analyzed genotypes; thus, the content of photosynthetic pigments was measured and shown in (**e**). (**f**) Expression of *CIA2* and *CIL* in the analyzed genotypes was measured using qRT-PCR with gene-specific primers. Locations of primers are depicted in panel (**a**). (**g**, **h**) Complementation of *cia2–2* phenotype with ectopic expression of *CIA2* under 35S and native promoter. (**g**) Relative expression of *CIA2* in analyzed lines. (**h**) Relative content of photosynthetic pigments in complementation lines. In (**a**), (**e**) and (**f**) statistical significance (ANOVA and Tukey HSD test) is shown relative to Col-0 (\*\**p* < 0.01; \*\*\**p* < 0.001). In (**g**) and (**h**) statistical significance (ANOVA and Tukey HSD test) is shown relative to *cia2-2* (\*\**p* < 0.01; \*\*\**p* < 0.001) and to Col-0 (^*p* < 0.05; ^^*p* < 0.01; ^^^*p* < 0.001).

### CIA2 and CIL function redundantly

To confirm the role of *CIA2* under the analyzed stress conditions, we isolated two additional independent *cia2* mutant alleles, *cia2–2* (SALK_004037, Col-0 background) and *cia2–3* (SGT49, Ler-0 background). In both mutants, the T-DNA was inserted into the first exon of *CIA2* (Figure 1c). Because *cia2–4* is an intronic allele, we decided to focus our analysis on *cia2–2* and *cia2–3* alleles. To test whether the incorporation of T-DNA into the *CIA2* gene inhibits the accumulation of *CIA2* transcript, *cia2–2* and *cia2–3* mutants were subjected to quantitative RT-PCR analysis using two pairs of primers (Figure 1c,d and Figure S3d). Although we did not detect *CIA2* transcript in *cia2–2* using primers flanking the T-DNA insertion site, we detected the PCR product (although significantly lower than that in Col-0 and *cil-1*) using primers specific to 3’UTR of the *CIA2* gene suggesting the presence of truncated CIA2 transcript in *cia2–2* (Figure 1d). In *cia2–3*, we did not detect PCR products in both tested primer pairs (Figure S3d).

CIA2 shares 54% identical amino acids with CIA2-LIKE (CIL, AT4G25990); moreover, CIL also contains C-terminal CCT motif, N-terminal cTP and NLS (Figure 1c). Because of this sequence similarity, CIL was speculated to act redundantly to CIA2 (Sun *et al.*, 2001). To test this hypothesis, we isolated *cil-1* (SAIL_228_C01, Col-0 background) and *cil-2* (SK14786, Col-4 background) mutants and introduced them into *cia2–2* background by crossing. Despite complete lack of *CIL* transcript in *cil-1* and *cil-2* mutants (Figure 1d and Figure S3a), both mutants were phenotypically indistinguishable from their corresponding wild-type (WT) plants. However, the double mutants *cia2–2 cil-1* and *cia2–2 cil-2* were paler than WT and the corresponding single mutants (Figure 1e and Figure S3b). To confirm visual differences, we measured the concentrations of photosynthetic pigments (Sumanta *et al.*, 2014). From this analysis, we deduced that the content of chlorophyll *a*, chlorophyll *b*, and total carotenoids is significantly lower in *cia2–2* and *cia2–3* (Figure 1f and Figure S3) mutants, thus supporting previous results obtained for *cia2–1* mutant (C.,-W., Sun *et al.*, 2009). The introduction of *cil* mutations into *cia2–2* background further reduced the concentration of photosynthetic pigments (Figure 1e,f and Figure S3b,c), suggesting that CIA2 and CIL act in an unequally redundant manner.

To further confirm that the lack of CIA2 is responsible for the observed phenotype, we introduced constructs that encode CIA2:YFP and YFP:CIA2 fusion proteins under CIA2 native promoter and CIA2:YFP under 35S promoter into *cia2–2* background. The complementation lines p*CIA2∷CIA2:YFP*_*cia2–2*_ and p*CIA2∷YFP*_*cia2–2*_ exhibited *CIA2* transcript level comparable to that of WT plants, whereas a ~7-fold increase was observed in p35S∷CIA2:YFP plants (Figure 1g). The complementation and overexpression lines were visually indistinguishable from those of WT plants, and spectrophotometric measurement confirmed that the obtained transgenic lines had WT-like levels of photosynthetic pigments (Figure 1h). However, despite many efforts, we were unable to detect the YFP signal in these complementation lines, which might be related to the high proteolytic turnover rate of CIA2. Taken together, our results strongly suggest that CIA2 and CIL act redundantly and are required for the normal accumulation of photosynthetic pigments and thus for the proper functioning of photosystems and photosynthesis.

### Characterization of *cia2 cil* responses in chloroplast-targeted stress

Because *cia2–4* mutant was initially selected from the screening using stress conditions that target the photosynthetic apparatus (Figure 1a,b), we set out to test the importance of *CIL* and *CIA2* under stress conditions using chlorophyll fluorescence as a readout. c*ia2–4* was shown to be more susceptible to UV-AB that induced cell death (measured with ion leakage and chlorophyll fluorescence) to a significantly higher extent than in Col-0 plants (Figure 1a). In agreement with these data, the exposure of *cia2–2* mutant to UV-AB resulted in similar changes in the photosynthetic parameters, which were significantly affected 1 and 2 days after the treatment (Figure 2a-c). Moreover, the introduction of *cil* mutant into *cia2–2* intensified the effect (Figure 2 and Figure S4). Measurements of ion leakage after UV-AB treatment confirmed chlorophyll fluorescence observations suggesting increased cell death in *cia2–2* and even stronger effect in *cia2–2 cil-1* and *cia2–2 cil-2* (Figure 2 and Figure S4).

**Figure 2.**
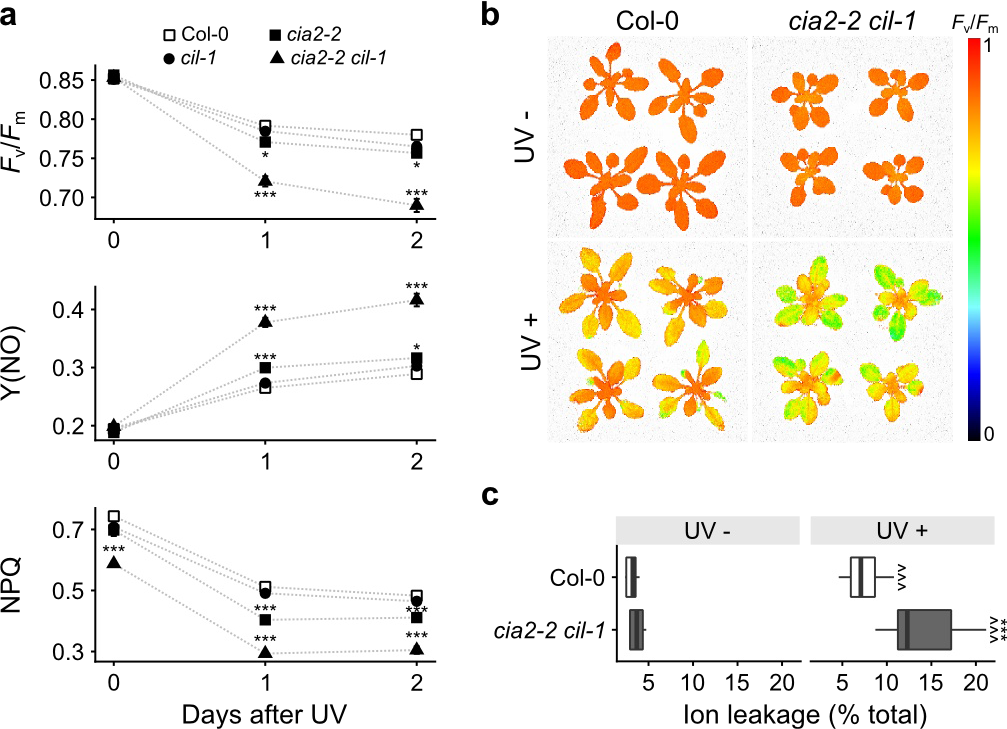
CIA2 and CIL are required for resistance to UV-AB. Mature plants were exposed to UV-AB, and the plants’ performance was assessed using chlorophyll *a* fluorescence (**a** and **b**) and ion leakage (**c**). **(a)** Maximum efficiency of PSII (*F*_v_/*F*_m_), nonregulated energy dissipation (Y(NO)), and nonphotochemical quenching (NPQ) were measured before and after UV-AB treatment (1 and 2 days). Each point represents mean ±SEM of at least eight plants. (**b**) *F*_v_/*F*_m_ of control (UV-) and UV-AB-treated (UV +) plants of Col-0 and *cia2–2 cil-1* plants. Values of *F*_v_/*F*_m_ are shown in pseudocolor scale. **(c)** Ion leakage of control (UV-, n = 5) and UV-AB-treated (UV +, n = 18) plants is shown as a percentage of total ion leakage. Statistical significance (ANOVA and Tukey HSD test) is shown relative to Col-0 (\**p* < 0.05; \*\*\**p* < 0.001) and to control conditions (^^^*p* < 0.001).

To confirm the role of CIA2, and test whether CIL is required for the high light response, we exposed the plants to blue HL (bHL) and monitored *F*_v_/*F*_m_ (Figure 3 and Figure S4). At all tested time points, we observed stronger photoinhibition of PSII as evidenced by decreased *F*v/*F*m in *cia2–2 cil-1* and *cia2–2 cil-2* (compared to that in WT plants) measured at a level of whole rosette with the strongest effect visible in young (not fully developed) leaves (Figure 3a,b and Figure S4e). As increased photoinhibition can result from either increased damage of PSII or decreased PSII repair (Miyata *et al.*, 2015), we monitored recovery after exposure to bHL (Figure 3c). We did not observe differences between Col-0 and *cia2–2 cil-1* in the rate of *F*_v_/*F*_m_ recovery suggesting that increased susceptibility to bHL in *cia2–2 cil-1* is not related to the repair of PSII but to increased damage. HL-induced damage of the photosynthetic electron transport chain is often related to the production of ROS such as singlet oxygen, which is mainly produced in PSII antenna, and superoxide anion and H_2_O_2_, which are mainly produced in the vicinity of PSI (Smirnoff and Arnaud, 2019). To simulate conditions of photoinhibition at the PSII side and induce overproduction of ROS, we treated leaf disks of Col-0 and *cia2–2 cil-1* with 3-(3,4-dichlorophenyl)-1,1-dimethylurea (DCMU) and methyl viologen (MV) and monitored *F*_v_/*F*_m_ (Figure 3d). DCMU noncovalently binds to the quinone B binding side of PSII and inhibits the photosynthetic electron transfer to the plastoquinone (PQ) pool, thus keeping reduced quinone A and oxidized PQ pool. MV is able to induce the accumulation of superoxide anion in chloroplast stroma. Upon DCMU and MV treatments, *F*_v_/*F*_m_ significantly more decreased in *cia2–2 cil-1* than in Col-0, suggesting that the double mutant is more susceptible to photoinhibition and ROS such as singlet oxygen and H_2_O_2_ (Figure 3d). Collectively, our results suggest that CIA2 and CIL are required for an adequate response to conditions that promote photooxidative stress in chloroplasts and trigger cell death signaling.

**Figure 3.**
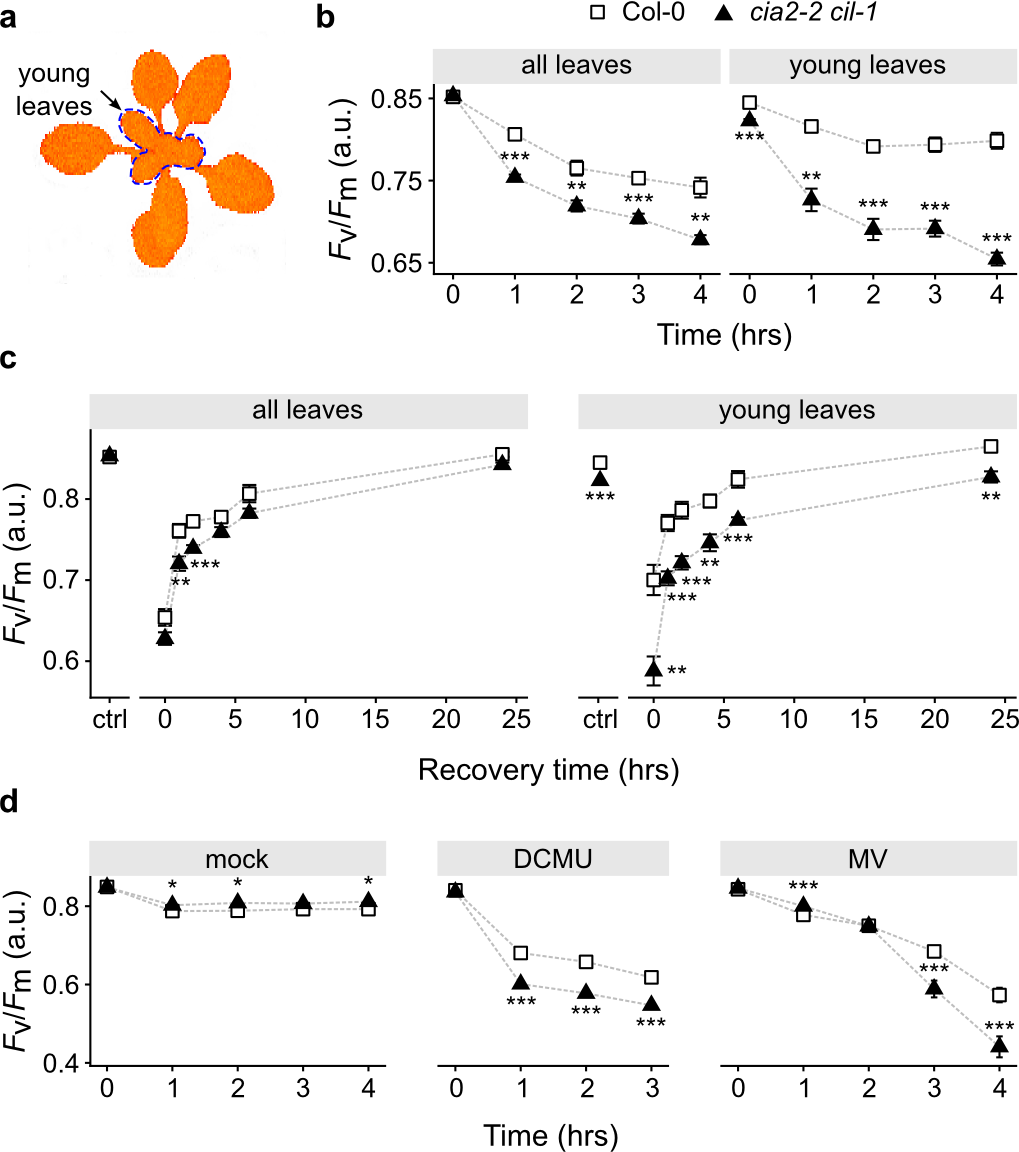
High-light (HL) susceptibility of *cia2–2 cil-1*. (**a**) Representation of Arabidopsis rosette with the young leaves marked with blue, dashed line. (**b, c, d**) Maximum efficiency of PSII (*F*_v_/*F*_m_) measured in plants (**b**) exposed to blue HL (1100 μmol m^−2^ s^−1^) for a specific time (n = 4–8 plants), (**c**) during recovery after HL stress, n = 6–8, (**d**) and treated with inhibitors influencing the production of ROS in chloroplasts (n = 12–22). In each plot, points represent mean ±SEM measured in independent plants (**b** and **c**) or leaf disks (**d**). Statistical significance (ANOVA and Tukey HSD test) is shown relative to Col-0 (\**p* < 0.05; ** *p* < 0.01;\*\*\**p* < 0.001)

### Role of CIA2 and CIL in the regulation of photosynthesis and NPQ

Increased susceptibility to HL and UV-AB can be a consequence of impaired nonphotochemical quenching (NPQ), which is responsible for the dissipation of excess light energy as heat (Kulasek *et al.*, 2016; Białasek *et al.*, 2017). Thus, we monitored NPQ using chlorophyll *a* fluorescence in Col-0, *cia2–2*, *cil-1*, *cil-2*, *cia2–2 cil-1*, and *cia2–2 cil-2* (Figure 4 and Figure S5). We observed that NPQ was slightly (but statistically significant) decreased in *cia2–2 cil-1* and *cia2–2 cil-2* at a level of the whole rosette. Differences were more pronounced when only young, not fully developed leaves were analyzed. In young leaves of *cia2–2 cil-1*, NPQ was decreased by 40% when compared to Col-0 (Figure 4a,b). It is well established that the process of NPQ is controlled by a small PSII protein PsbS, which is activated by the acidification of the thylakoid lumen (Niyogi *et al.*, 2005). To check whether this process is impaired in *cia2 cil* mutants, we measured the proton gradient (ΔpH) and electric potential (Δψ), which constitute proton motive force (PMF) (Figure 4c-e and Figure S5c-e) at two actinic light intensities (i.e., 160 and 660 μmol m^−2^ s^−1^). Although we did not observe differences in PMF between the analyzed genotypes, we noted that *Δp*H in *cia2–2* as well as in *cia2–2 cil-1* and *cia2–2 cil-2* was significantly decreased as compared to that in Col-0 (Figure 4e and Figure S5e), which correlates well with decreased NPQ in these genotypes. Because the rosette size of *cia2–2 cil-1* was slightly smaller than that of Col-0 (Figure 1d, Figure 4b), we also measured CO_2_ assimilation as a function of light intensity (20–2000 μmol m^−2^ s^−1^) and CO_2_ concentration (20–1500 ppm) (Figure 4f,g) and observed moderate but statistically significant decrease in CO_2_ assimilation in *cia2–2 cil-1* within the tested range of light intensity and CO_2_ concentration. Taken together, our results strongly suggest that CIA2 and CIL are required for optimal NPQ and proper formation of trans-thylakoid proton gradient as well as for optimal CO_2_ assimilation.

**Figure 4.**
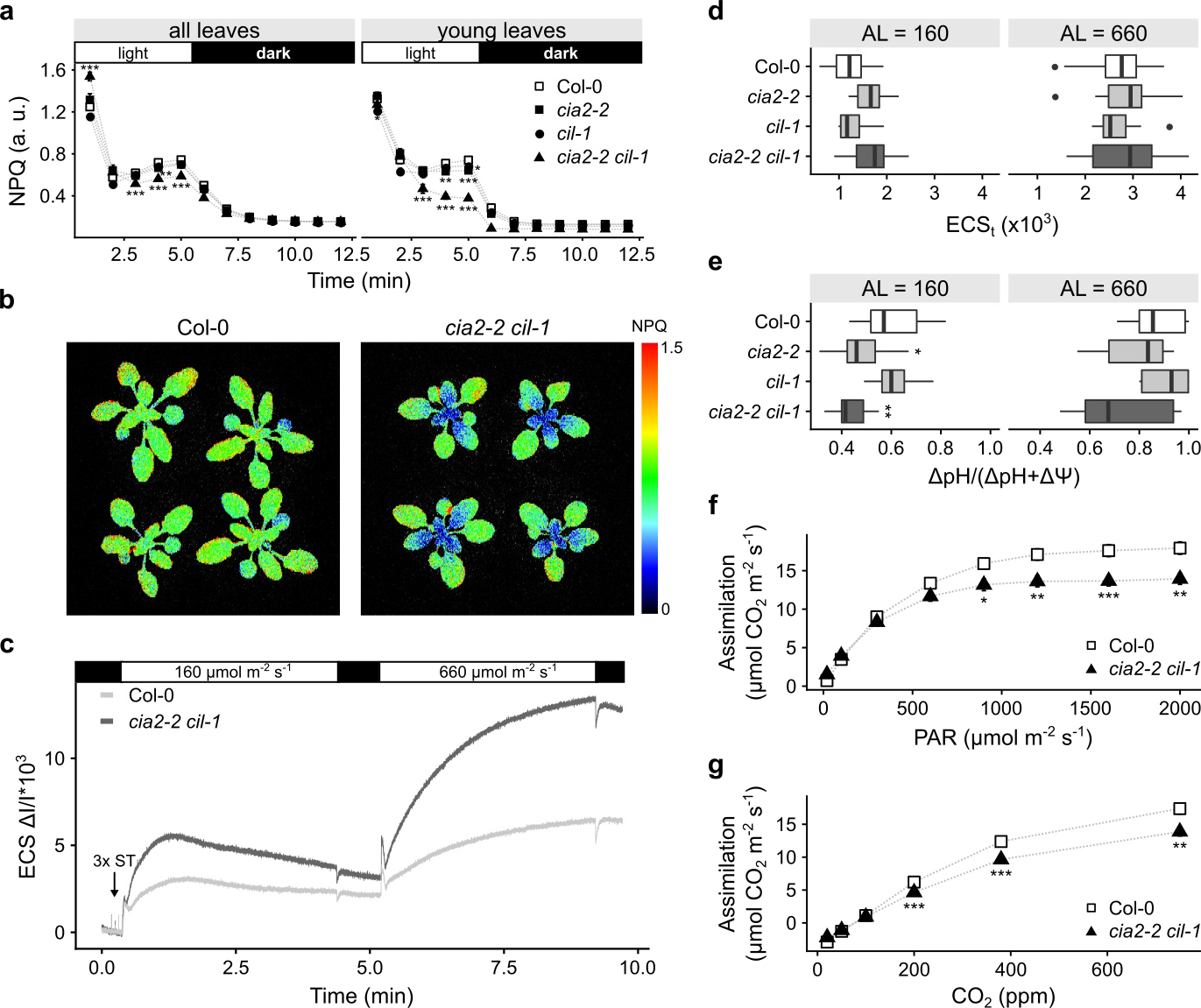
CIA2 and CIL are required for optimal photosynthesis in Arabidopsis. (**a**) Nonphotochemical quenching (NPQ) in analyzed genotypes. Points represent mean ±SEM of the whole rosette and only young leaves. (**b**) NPQ of Col-0 and *cia2–2 cil-1*. (**c**) Analysis of electrochromic pigment shift (ECS, P515) at 160 and 660 μmol m^−2^ s^−1.^ of actinic light. For simplicity, only Col-0 and *cia2–2 cil-1* are shown. ST–single turnover flash. Total ECS (ECSt) (**d**) and ΔpH (**e**) in analyzed genotypes at 160 and 660 μmol m^−2^ s^−1^. Box plots represent values of 10 independent plants. (**f**) CO_2_ assimilation as a function of light intensity. (**g**) CO_2_ assimilation as a function of CO_2_ concentration. In (**f**) and (**g**), values represent mean ±SEM of 7–9 plants. Statistical significance (ANOVA and Tukey HSD test) is shown relative to Col-0 (\**p* < 0.05; \*\**p* < 0.01; \*\*\**p* < 0.001)

### Chloroplast translation is attenuated in *cia2 cil*

It was shown that CIA2 binds to the promoters of genes encoding the chloroplast ribosomal proteins, and decreased expression of these genes was observed in *cia2–1* mutant (Sun *et al.*, 2001). To check whether a similar phenomenon was observed in *cia2–2* and if the expression of these genes was dependent on the activity of CIL, we checked the expression of *Rps6*, *Rpl11*, *Rpl18*, and *Rpl28* nuclear genes encoding for bS6c, uL11c, uL18c, and bL28c chloroplast ribosomal proteins using qRT-PCR (Figure 5a). The expression of all analyzed genes was decreased in *cia2–2* as compared to that in Col-0 and *cil-1*. The introduction of *cil-1* mutation into *cia2–2* led to stronger reduction in the expression of all genes except *Rpl18*, whose transcript was at the same level as that in *cia2–2* (Figure 5a). To gain more comprehensive view of gene expression changes, we performed whole transcriptome sequencing (RNA-seq) for Col-0, *cia2–2*, *cil-1*, and *cia2–2 cil-1* plants. First, we focused on genes encoding for proteins involved in chloroplast translation. Our analysis revealed that the expression of 21 out of 66 chloroplast ribosome genes is inhibited in *cia2–2 cil-1* (Figure 5b) including two genes encoded by the chloroplast genome (i.e., *rpl2* and *rpl32*, encoding for the ribosomal proteins uL2c and bL32c). Pentatricopeptide repeat (PPR) proteins are involved in every step of chloroplast gene expression, namely transcription, RNA metabolism, and translation (Barkan and Small 2014). Thus, we checked the expression of chloroplast-targeted PPRs in *cia2–2 cil-1* and found 13 down-and 7 upregulated PPR genes (Figure 5b). Further, the expression analysis of other genes possibly involved in plastid translation revealed the presence of both up-and downregulated genes (Figure 5b). Taken together, the transcriptome profiling indicates that the lack of *CIA2* and *CIL* leads to altered expression of multiple genes encoding the components of the chloroplast translation machinery.

**Figure 5.**
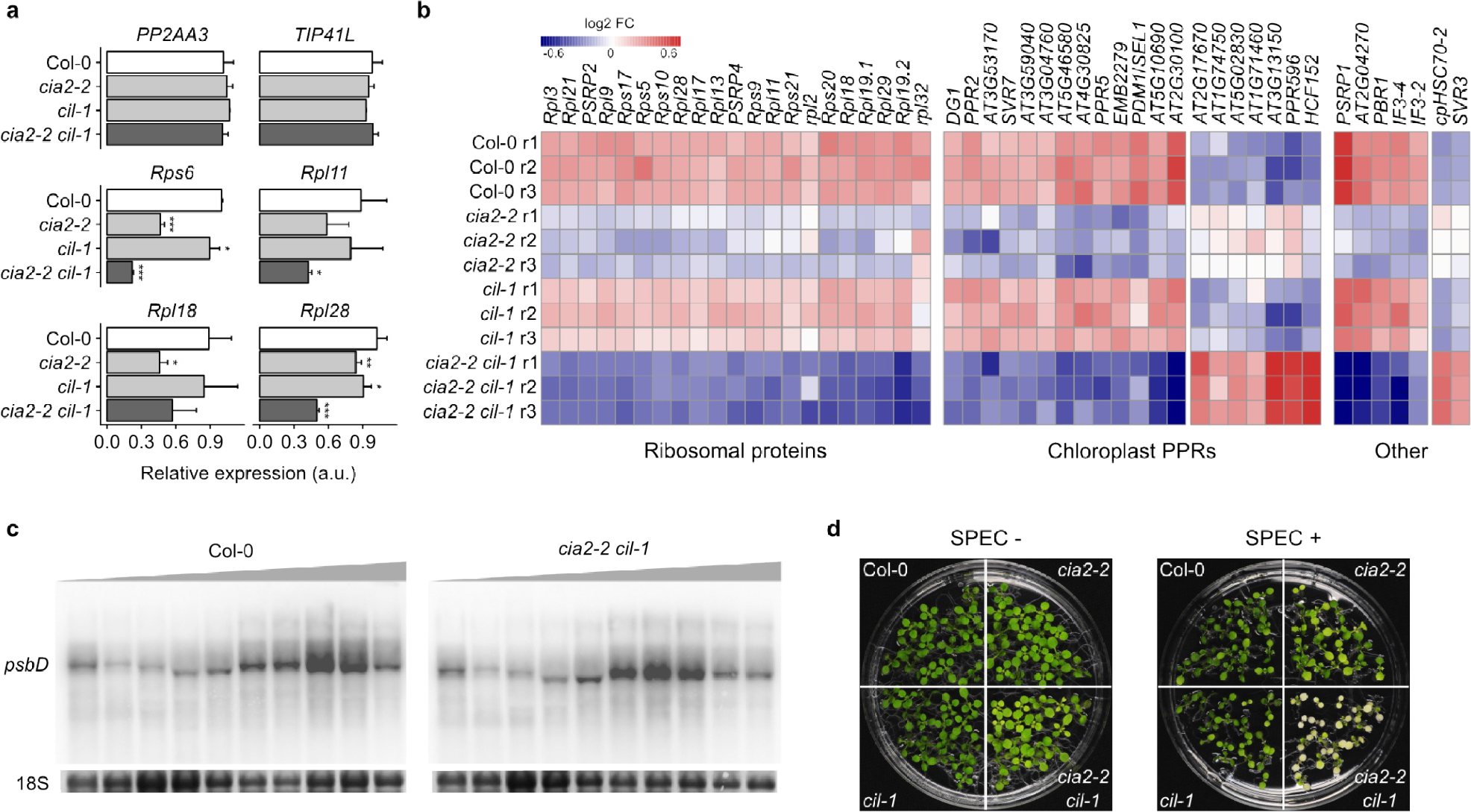
Characterization of chloroplast translation in analyzed genotypes. (a) Expression analysis of genes encoding chloroplast ribosomal proteins. Levels of analyzed transcripts were determined using qRT-PCR and normalized to two house-keeping genes (*PP2AA3* and *TIP41L*). Bars indicate mean values±SD (three independent biological replicates). (b) Expression profile (RNA-seq) of genes involved in the regulation of plastid translation. (c) Polysome analysis reveals slower translation of *psbD* mRNA in *cia2–2 cil-1* double mutant as compared to Col-0. Gray triangles indicate the density of sucrose gradient. Methylene blue-stained 18S rRNA is shown as a loading control. (d) Susceptibility to chloroplast translation inhibitor (spectinomycin, 1.25 mg/L) of analyzed genotypes. Statistical significance (ANOVA and Tukey HSD test) is shown relative to Col-0 (\**p* < 0.05; \*\**p* < 0.01; \*\*\**p* < 0.001).

To further confirm that CIA2 and CIL are required for optimal chloroplast translation, we subjected Col-0 and *cia2–2 cil-1* to polysome loading assays. We examined the ribosomal loading of *psbD* mRNA (encoding for the D2 protein of PSII) using sucrose gradient fractionation followed by northern blot analysis (Figure 5c). This analysis showed that *psbD* mRNA was underrepresented in the polysomal fractions in *cia2–2 cil-1* compared to that in Col-0, and the mRNA was shifted toward lighter (monosome) fractions, suggesting that *psbD* mRNA was associated with fewer ribosomes and that its translation was reduced in *cia2–2 cil-1*. Further, as multiple lines of evidence indicate that CIA2 and CIL are involved in chloroplast translation, we also tested the growth of Col-0, *cia2–2*, *cil-1*, and *cia2–2 cil-1* in media supplemented with spectinomycin, which inhibits this process. Spectinomycin binds the 30S subunit of the 70S ribosome and prevents translocation of peptidyl-tRNA from A to P site; consequently, many Arabidopsis translation mutants showed increased susceptibility to this antibiotic (Parker *et al.*, 2014). In agreement with the results of ribosome loading experiments, we observed severe inhibition of pigment accumulation in *cia2–2 cil-1* grown in media supplemented with spectinomycin (Figure 5d), which suggests that CIA2 and CIL play a role in plastid translation.

The decreased chlorophyll content in young leaves and inhibited translation in *cia2–2 cil-1* led us to analyze chloroplast ribosomal RNA (rRNA) maturation. In Arabidopsis chloroplasts, all rRNAs are encoded by the polycistronic *rrn* operon (Bollenbach *et al.*, 2007). Upon processing of the initial transcript by distinct endo-and exonucleases, tRNAs, precursors of 16S and 5S rRNAs and bicistronic 23S-4.5S intermediate are created (Figure 6a). The 23S-4.5S precursor is cut to produce 4.5S and 23S rRNA fragments. The maturation of 23S rRNA is followed by the introduction of two gaps (“hidden breaks”) producing three distinct parts of 0.4, 1.1, and 1.3 kb; however, the functional relevance of this postmaturation processing is not clear (Bollenbach *et al.*, 2007; Fristedt *et al.*, 2014). The capillary electrophoresis of RNA isolated from young leaves showed that 1.1 and 1.3 kb species of 23S rRNA and 16S rRNA accumulate to lower level in *cia2–2* and *cia2–2 cil-1* mutants as compared to that in WT plants (Figure 6b). These results suggest that the expression and/or processing of 16S and 23S rRNAs are dependent on CIA2 and CIL. To further characterize rRNA maturation, we performed northern blot analysis with probes specific to *rrn16* and *rrn23* (Figure 6c). We observed only slightly lower level of 16S rRNA in *cia2–2 cil-1* without accumulation of its 1.7 kb precursor suggesting that the processing of *rrn16* is not impaired (Figure 6c). In agreement with the results of capillary electrophoresis, we observed decreased accumulation of 0.4, 1.1, 1.3, and 1.7 kb species and increased accumulation of 2.4 kb fragment of 23S rRNA in *cia2–2 cil-1* (Figure 6c). These results suggest that gap incorporation between 1.1 kb and 1.3 kb fragments is less efficient in *cia2–2 cil-1* mutant. To confirm these results, we performed qRT-PCR analysis with primers spanning hidden breaks (Figure 6d) and obtained confirmatory results.

**Figure 6.**
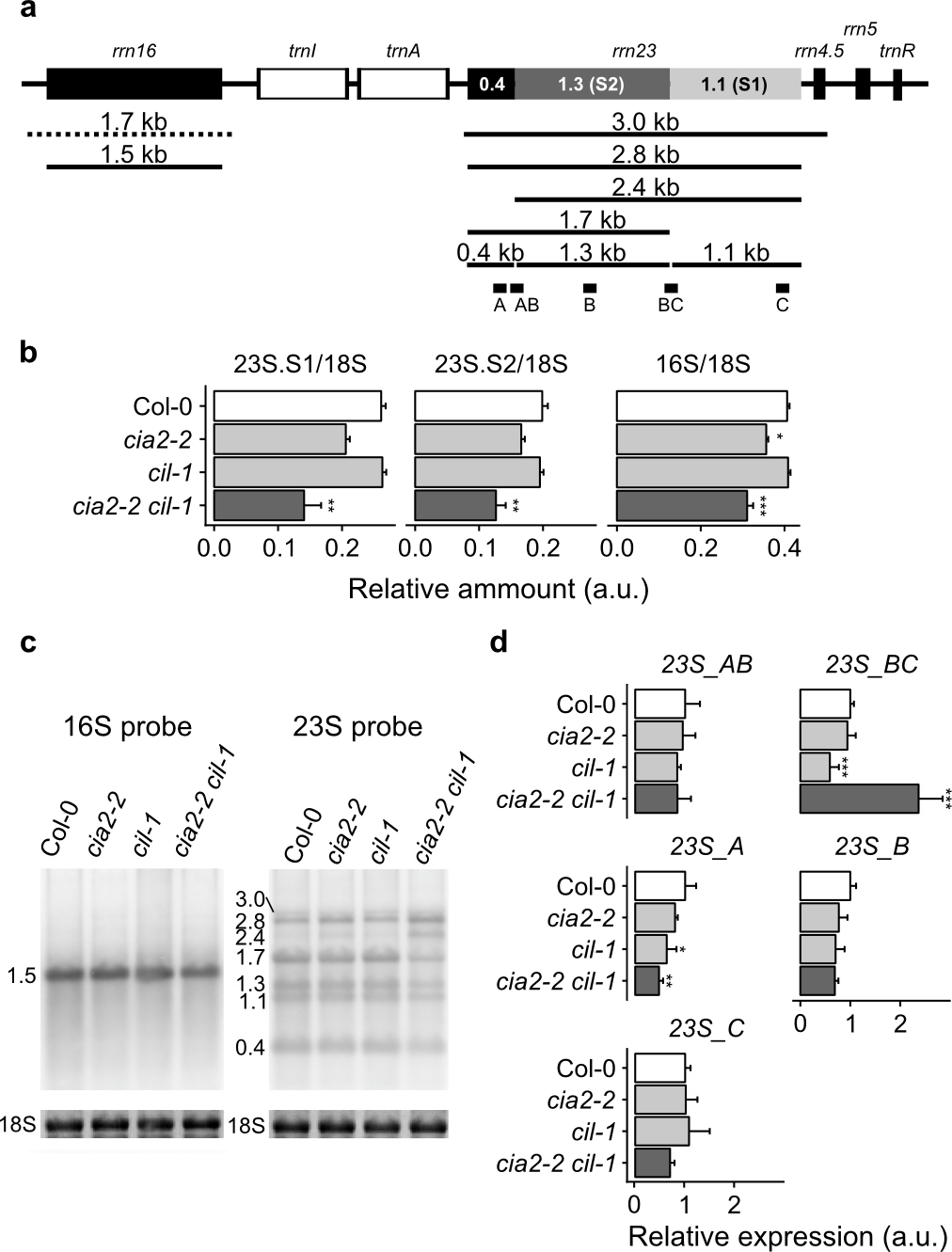
Maturation of plastid rRNAs. (**a**) Plastid rRNA operon. Mature forms of 16 and 23S rRNA are shown. Precursor of 23S rRNA is processed and cleaved into three parts: 0.4, 1.1, and 1.3 kb. (**b**) Relative abundance of precursor 23S and 16S rRNAs was measured using capillary electrophoresis and normalized to the cytoplasmic 18S rRNA. Values represent mean values ± SEM (n = 3). (**c**) Maturation and abundance of plastid rRNA were measured using Northern blot indicating that the 2.4 kb form of 23S rRNA is accumulated in *cia2–2 cil-1*. (**d**) Determination of relative amounts of 23S rRNA forms with qRT-PCR and primers (depicted in panel (**a**)) specific to each of cleaved fragments (primers A, B, and C) and flanking “hidden breaks” (primers AB and BC). Values were normalized to the level of 16S rRNA and represent mean values ± SD (n = 3). Statistical significance (ANOVA and Tukey HSD test) is shown relative to Col-0 (\**p* < 0.05; \*\**p* < 0.01; \*\*\**p* < 0.001).

Next, we tested whether the perturbations of chloroplast translation are linked to the decreased NPQ observed in *cia2 cil* (Figures 4f and S6). For this, we measured NPQ in well-characterized chloroplast translation mutants *prpl11–1*, *prps1–1*, *psrp5-R1*, and *rps17–1* (Pesaresi *et al.*, 2001; Romani *et al.*, 2012; Tiller *et al.*, 2012; Tadini *et al.*, 2016) (Figure S6). However, in contrast to *cia2 cil*, NPQ in these mutants was increased as compared to that in Col-0 plants (Figure S6). These results suggest that the observed decrease in NPQ in *cia2 cil* mutant is not related to inhibited chloroplast translation. Taken together, our results suggest that the translation of chloroplast mRNA and the maturation and accumulation of 23S rRNA are influenced by CIA2 and CIL.

### Lack of CIA2 and CIL confers heat stress tolerance

To our surprise, RNA-seq analysis showed that many genes induced in *cia2–2 cil-1* were annotated as heat shock proteins (HSPs) (Data S2). Further, the gene ontology (GO) enrichment analysis demonstrated that GO terms related to heat response and acclimation were overrepresented among genes induced in *cia2–2 cil-1* (Figure 7a). To confirm the RNA-seq results, we performed qRT-PCR analysis to assess the transcript level of *HEAT SHOCK TRANSCRIPTION FACTOR A2 (HSFA2)* and *HEAT SHOCK PROTEIN 70–4* (*HSP70–4*) in *cia2–2, cia2–3*, *cil-1*, *cia2–2 cil-1*, and corresponding WT plants (Figure 7b). Our results confirmed strong induction of both genes in *cia2* mutants, and even stronger induction was observed in *cia2–2 cil-1* double mutant (Figure 7b). To check whether increased expression of many HSPs can confer thermotolerance, we grew *cia2–2, cil-1, cia2–2 cil-1*, and Col-0 seedlings on Petri dishes and exposed them to heat stress (45 ^o^C) for 20–40 min (Figure 7c,d). Exposure to 45 °C for 30 min resulted in almost complete bleaching and growth inhibition of Col-0 plants (Figure 7c). A similar phenotype was observed in the case of *cia2–2* and *cil-1* mutants; however, *cia2–2* c*il-1* plants were affected to a significantly lower extent, and nearly all individuals retained cotyledon and leaf pigmentation (Figure 7c,d). Furthermore, at other tested time points, *cia2–2 cil-1* was more resistant to heat stress than the remaining genotypes (Figure 7c,d). These results suggest that CIA2 and CIL are redundantly involved in the modulation of thermotolerance presumably through the regulation of HSP expression.

**Figure 7.**
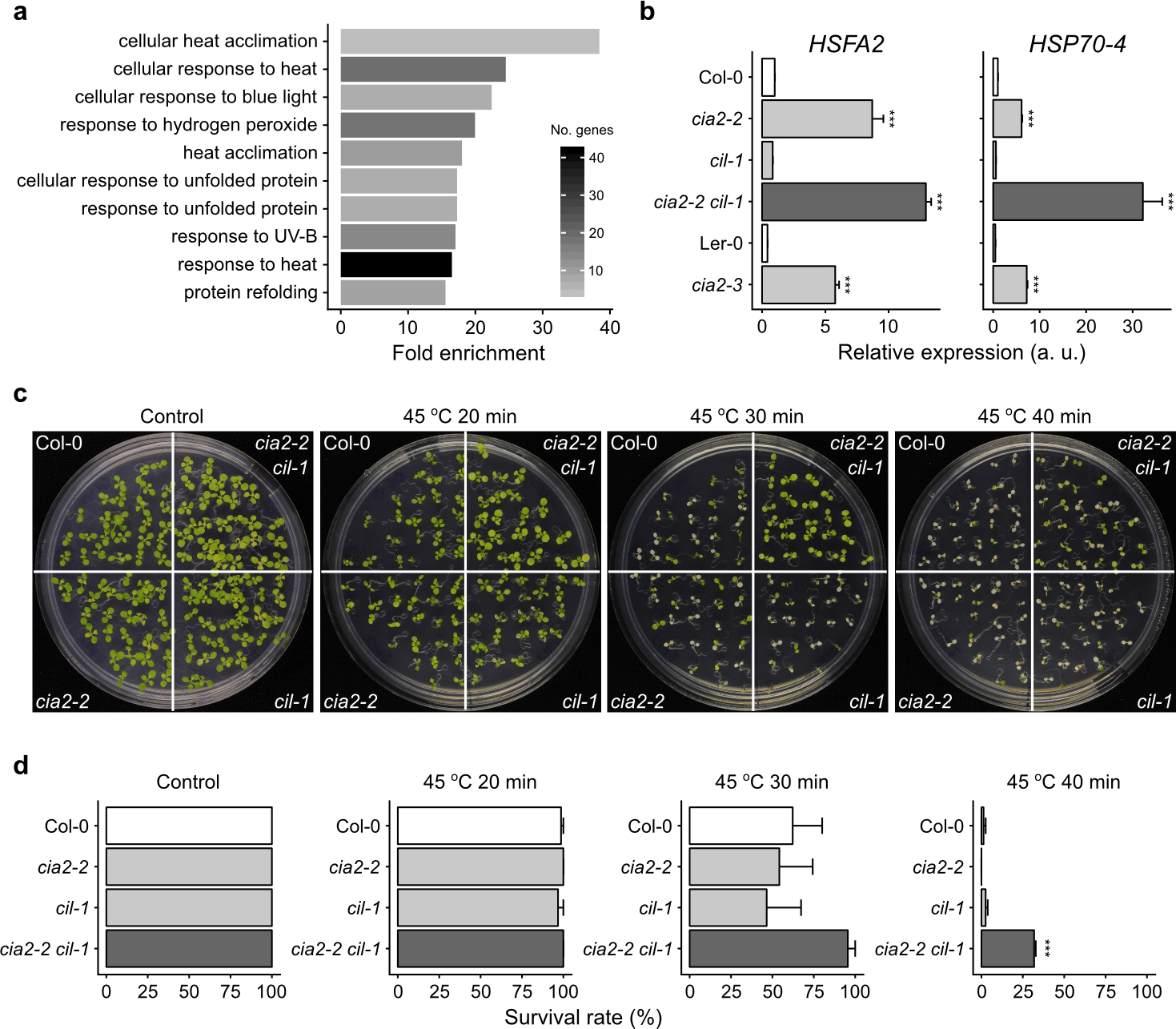
CIA2 and CIL negatively regulate tolerance to heat shock. (**a**) Gene ontology (GO) analysis of genes significantly induced in *cia2–2 cil-1* compared to Col-0 in the RNA-seq experiment. Ten most significantly overrepresented GO terms are shown. Number of genes in each group is shown in color of the bar. (**b**) Validation of expression of heat shock marker genes using qRT-PCR in analyzed genotypes and additional allele (*cia2–3*) in Ler-0 background (n = 3). Error bars represent ± SD. (**c** and **d**) Thermo tolerance was tested in seedlings grown *in vitro*. Plants were exposed to 45 °C for a specific period of time. (c) Pictures of plants grown in control conditions 7 days after heat shock. (**d**) Survival rate of control and heat shock-treated plants. Error bars represent ±SEM (n ≥ 3 plates) from two independent experiments (in total at least 75 seedlings were analyzed per genotype and treatment). Statistical significance (ANOVA and Tukey HSD test) is shown relative to corresponding WT (\*\*\**p* < 0.001).

## Discussion

Mature chloroplasts contain approximately 3000 proteins, the majority of which are located in the stroma and thylakoid membranes (Jarvis and Robinson, 2004). Chloroplasts are capable of semiautonomous protein synthesis; however, their genomes encode only approximately 80 proteins. More than 95% of their proteins are products of the nuclear genome, which are translated in the cytosol and then imported (Jarvis and Robinson, 2004). Plastid biogenesis and maintenance depend on the coordinated assembly of proteins imported from the cytosol with proteins translated within plastids (Jarvis and López-Juez, 2013). Chloroplasts in leaf cells have a greater need for protein import and protein synthesis than plastids in other organs due to a large number of proteins required for photosynthesis (Jarvis and Robinson, 2004). Nuclear-encoded proteins are transported across the double chloroplast membrane due to the activity of a specialized translocon complex (Nakai, 2015; Paila *et al.*, 2015). This mechanism is essential for chloroplast biogenesis and requires coordinated action of multiple proteins. Previous reports revealed a positive regulatory effect of CIA2 on the
transcription of the translocon genes Toc33 and Toc75 in leaves (Sun *et al.*, 2001; C.,-W., Sun *et al.*, 2009), suggesting the role of CIA2 in protein import into the chloroplast.

In the present study, we found that CIA2 has a functional cTP which, in transient expression experiments, targets a fraction of CIA2:YFP fusion protein to chloroplasts of Arabidopsis seedlings (Figure 1c, Figure S2). In agreement with our observation, recently characterized barley (*Hordeum vulgare*) CIA2 homolog, *Hv*CMF7, was also shown to be localized in chloroplasts (Li *et al.*, 2019). In the first attempt to determine the subcellular localization of CIA2, the GUS:CIA2 fusion protein was found to reside exclusively in the nucleus (Sun *et al.*, 2001). We speculate that the discrepancy between our data and the previous work might be related to the masking of CIA2 cTP by the GUS fused to CIA2 N-terminus, thus inhibiting the import of GUS:CIA2 fusion protein into chloroplasts (Sun *et al.*, 2001). From the results of our complementation test, we were unable to determine the significance of CIA2 chloroplast localization because the expression of both constructs, p*CIA2∷CIA2:YFP* and p*CIA2∷YFP:CIA2*, restored the accumulation of photosynthetic pigments in *cia2–2* mutant (Figure 1h). Thus, we speculate that at least with regard to the determination of the levels of photosynthetic pigments, the chloroplast localization of CIA2 is not crucial. Interestingly, despite clear complementation effect and restoration of *CIA2* transcript level, we did not observe YFP signal in stable transformants expressing CIA2:YFP fusion either under native or 35S promoter, suggesting that the CIA2 protein level might be subject to strict posttranslational control. A similar scenario was observed for ABI4 (Finkelstein *et al.*, 2011) and RCD1 (Jaspers *et al.*, 2009), suggesting that certain regulatory proteins might be unstable and/or degraded by the proteasome thus preventing their detection in standard conditions. A comparable phenomenon was observed in the case of another key player of retrograde signaling, GUN1, which accumulates in chloroplasts only at a very early stage of leaf development (Wu *et al.*, 2018).

Inefficient chloroplast protein import causes cytosolic overaccumulation of preproteins, which results in the activation of chaperones such as HSP70 and HSP90, which were recently postulated to be key components of the chloroplast retrograde signaling pathway (Wu *et al.*, 2019). Accordingly, we observed strong transcriptional induction of cytosolic chaperones and increased thermotolerance in *cia2 cil* mutant (Figure 7). On the other hand, it was demonstrated that thermotolerance is regulated by chloroplast signals that depend on the redox state of the PQ pool or hydrogen peroxide produced in chloroplasts (Dickinson *et al.*, 2018).

The redox state of the PQ pool and H_2_O_2_ are also involved in high-light acclimatory responses (Karpiński *et al.*, 1999; Mühlenbock *et al.*, 2008; Gilroy *et al.*, 2016). Experimentally, it is almost impossible to separate foliar heat shock from high-light responses, because exposure to high light for a few seconds significantly warms up Arabidopsis leaves due to the dissipation of energy as heat (Kulasek *et al.*, 2016). Therefore, it is interesting that CIA2 and CIL antagonistically influence high-light and UV-B acclimation *versus* thermotolerance in ambient light. Abolished expression of *CIA2* and *CIL* genes in double mutant causes growth reduction, acclimation, and photosynthesis dysfunction, while it can cause thermotolerance expressed as seedling survival rate at high temperatures. However, we were not able to identify the precise role of CIA2 and CIL in the antagonistic regulation of these processes, and we plan to perform field experiments similar to those we did for cell death conditional regulators (i.e. LSD1, EDS1, PAD4) (Wituszyńska *et al.*, 2013). These proteins, depending on growing conditions, differently regulate chloroplast retrograde cell death signaling for growth, photosynthesis, high-light and UV-B acclimation, water use efficiency, and seed yield (Wituszyńska *et al.*, 2013; Wituszyńska *et al.*, 2015; Bernacki *et al.*, 2019).

Our data show that CIA2 and its close homolog CIL play a relevant function in the chloroplast biogenesis process (Figure 1). The *cia2 cil* double mutant exhibited a pale green phenotype, which was much more pronounced than that observed in single *cia2* mutants, suggesting that these proteins act in an unequally redundant manner. Double mutant plants displayed a significantly lower concentration of chlorophyll *a*, chlorophyll *b*, and carotenoids (Figure 1 and Figure S3), suggesting impairment in the conversion of light into chemical energy. A similar phenotype was observed in plants lacking the activity of Golden 2-like (GLK) transcription factors. In the nuclear genome, GLKs are involved in the related expression of nuclear chloroplast-localized proteins and photosynthesis-related genes in *Zea mays*, *Physcomitrella patens*, and *Arabidopsis thaliana* (Yasumura *et al.*, 2005; Waters *et al.*, 2009). Like *cia2 cil*, the *glk1 glk2* double mutants were pale green and deficient in the formation of the photosynthetic apparatus (Waters *et al.*, 2009). Our RNA-seq and qRT-PCR-based expression data revealed a significant transcriptional downregulation of chloroplast ribosome genes in *cia2 cil* plants. Almost one-third of 66 chloroplast ribosome protein genes were significantly repressed in *cia2–2 cil-1*. These data are consistent with previous reports showing that CIA2 binds to the promoters of genes encoding chloroplast ribosomal proteins (C.,-W., Sun *et al.*, 2009) and support a positive regulatory role of CIA2 in chloroplast translation. Chloroplast ribosomal proteins are encoded by both nuclear and chloroplast genomes (Zoschke and Bock, 2018). Interestingly, the majority of ribosomal genes downregulated in *cia cil* are nuclear-encoded, which, together with partial chloroplast localization of CIA2 (Figure S2), suggests that CIA2 and CIL might act as chloroplast sensors of the environmental stimuli and mediate chloroplast-dependent adaptive responses (Estavillo *et al.*, 2013), and the dual chloroplast/nuclear localization of CIA2 (Figure S2) supports its involvement in this process.

Finally, our data indicate that CIA2 and CIL influence chloroplast translation by the regulation of ribosome assembly and maturation as *cia2 cil* double mutants displayed a disturbed accumulation of 23S rRNA (Figure 6). In plant chloroplasts, 23S rRNA constitutes a component of a large (50S) subunit of chloroplast ribosomes. The rRNA undergoes postmaturation processing, which includes a site-specific cleavage that generates gapped, discontinuous rRNA molecules (Nishimura *et al.*, 2010). The maturation process is followed by the removal of a specific region and introduction of a gap (the so-called hidden break) into the 23S rRNA (Bollenbach *et al.*, 2005). This molecule is split into four major fragments of 1.7, 1.3, 1.1, and 0.4 kb. Chloroplasts of all dicotyledonous plants have hidden breaks at similar locations. Several proteins have been reported to bind to a 23S rRNA segment close to the hidden break sites (Bieri *et al.*, 2017). We observed an increased accumulation of the 2.4 kb 23S rRNA species and a reduced occurrence of the hidden break in this species (Figure 6c,d). Interestingly, the expression of the two genes *Rpl19.1* and *Rpl19.2* (Figure 5b), which encode for the ribosomal protein bL19c binding close to the hidden break (Bieri *et al.*, 2017), was reduced. This indicates that the lack of bL19c could influence the rRNA structure in a way that makes it less susceptible to the hidden break incorporation. The lack of bL19c could be one of the causes of the impaired biogenesis of the 50S subunit as shown by the reduced accumulation of the 23S rRNA (Figure 6c,d). The deficiencies in rRNA maturation observed in *cia2–2 cil-1* double mutant suggest that CIA and CIL might play a role in this process, either directly by binding and processing 23S rRNA precursor or more likely indirectly by regulating the expression of ribosomal proteins.

In conclusion, the presented results demonstrate high complexity and elegancy of chloroplast retrograde signaling and anterograde nuclear responses that depend on putative TFs such as CIA2 and CIL. This complexity is not only observed on the molecular and biochemical levels but also on the physiological level of acclimatory responses as well as on the regulation of chloroplast biogenesis and plant growth and development. To better understand the interdependence of these regulatory processes, further interdisciplinary studies on the function of these proteins are needed.

## Experimental procedures

### Plant material and growth conditions

*Arabidopsis thaliana* ecotypes Columbia-0 (Col-0), Columbia-4 (Col-4), and Landsberg erecta (Ler-0) were used as controls for analyzed mutants. *cia2–2* (SALK_004037), *cia2–3* (SGT49), *cia2–4* (SALK_045340), *cil-1* (SAIL_228_C01), and *cil-2* (SK14786) were ordered from the Nottingham Arabidopsis Stock Centre (NASC) and genotyped using primers shown in Table S2. The *cia2–2 cil*-1 and *cia2–2 cil-2* double mutants were generated by crossing. Plants were grown on Jiffy pots (Jiffy Products) for 3–4 weeks in a growing chamber in long-day photoperiod (16 h/8 h) under constant white light of 140–160 μmol photons m^−2^ s^−1^ at 22 °C.

### Databases

In this work, we used information extracted from proteome databases to identify chloroplast proteins [Plant Proteomics Database at Cornell (http://ppdb.tc.cornell.edu/) (Q., Sun *et al.*, 2009); The Chloroplast Function Database II (http://rarge-v2.psc.riken.jp/chloroplast/) (Myouga *et al.*, 2013), and ARAMEMNON (http://aramemnon.uni-koeln.de) (Schwacke *et al.*, 2003)] and TF databases to look for annotated TFs in Arabidopsis [The Plant Transcription Factor Database (http://plntfdb.bio.uni-potsdam.de) (Pérez-Rodríguez *et al.*, 2009) and PlantTFDB (http://planttfdb.cbi.pku.edu.cn/) (Jin *et al.*, 2014)].

### Vector constructions and Arabidopsis transformation

*CIA2* and *CIL* full CDSs were amplified with primers listed in Table S2 by PCR using cDNA from Col-0 plants. PCR products of expected length were purified from agarose gel and inserted into the entry clone using the pENTR/D-TOPO cloning kit (Invitrogen). DNA fragments encoding for N-terminal 100 aa as well as predicted cTP were amplified from plasmids carrying *CIA2* and *CIL* CDSs. Next, they were subcloned into pGWB641 vector (Nakagawa *et al.*, 2007) using Gateway® LR Clonase™ II enzyme mix (Invitrogen). These constructs were introduced into *Agrobacterium fabrum* (GV3101). The vector was transformed into Col-0 and/or *cia2–2* mutant by the floral dip method. The T3 generation of homozygous transgenic plants was obtained and used for further experiments. The complementation lines were obtained in a similar manner, but in the final step, the promoter, CDS and YFP sequences were cloned into the pK7m34GW binary vector and transformed into Agrobacterium.

In parallel, whole *CIA2* and the *CIA2* fragment encoding for the 100 aa N-terminal part of *CIA2* were amplified from cDNA using USER compatible (Nour-Eldin *et al.*, 2006) primers and the improved Pfu X7 polymerase (Nørholm, 2010). PCR fragments were cloned into the pLIFE001 vector (Silvestro *et al.*, 2013) and introduced into Agrobacterium (GV3101) for transient expression in Arabidopsis seedlings and subcellular localization using the FAST method (J., F., Li *et al.*, 2009).

### Chlorophyll a fluorescence, CO_2_ assimilation, and ΔpH measurements

Chlorophyll *a* fluorescence parameters were measured in dark-acclimated (30 min) plants using PAM FluorCam 800 MF PSI device (Brno, Czech Republic) as described earlier (Gawronski *et al.*, 2014). CO_2_ assimilation was measured as described earlier (Burdiak *et al.*, 2015). The electrochromic pigment shift (ECS) was measured using DUAL-PAM 100 equipped with P515/535 module (Walz), which allows the simultaneous measurement of the dual beam 550–515 nm signal difference (Schreiber and Klughammer, 2008). Before measurement, the plants were dark acclimated for 30 min and subsequently illuminated with red actinic light of 160 μmol photons m^−2^ s^−1^ for 20 min. At the beginning of each measurement, the actinic light was turned off for 23 s, and three single turnover (ST) flashes were applied. Next, the red actinic light (160 μmol photons m^−2^ s^−1^) was turned on, and after 4 min, it was again turned off to determine total ECS, ΔpH, and ΔΨ as described previously (Herdean *et al.*, 2016). Immediately after the first measurement, the ECS signal was recorded using the same protocol but at a higher actinic light intensity (660 μmol photons m^−2^ s^−1^). The ECS signal was normalized to the highest ST peak recorded at the beginning of each measurement.

### Photosynthetic pigment analysis

For pigment analysis, approximately 15–20 mg of tissue was collected from the sixth leaf of 3–4-week-old plants. Tissue was immediately frozen in liquid nitrogen and ground in the presence of methanol. Absorbance was measured using Multiskan GO (Thermo Fisher Scientific) spectrophotometer after the clarification of the supernatant by centrifugation. Photosynthetic pigment concentrations were calculated as reported previously (Sumanta *et al.*, 2014).

### Stress treatments and ion leakage

High-light stress was applied to whole plants using blue light (455 nm) of 1100 μmol photons m^−2^ s^−1^ for the indicated time. As a light source, SL3500-B-D LED array was used (PSI, Brno, Czech Republic). After illumination, the plants were dark acclimated for 20 min, and chlorophyll a fluorescence was measured as described above. For recovery experiments, we used blue light of 2500 photons m^−2^ s^−1^ for 2.5 h. After this treatment, the plants were placed in the growing chamber for an indicated period of time followed by 20 min dark acclimation and chlorophyll *a* fluorescence measurement. For UV-AB stress, UVC 500 Crosslinker (Hoefer Pharmacia Biotech, USA) equipped with UV-A (TL8WBLB, Philips) and UV-B lamps (G8T5E, Sankyo Denki) was employed. The plants were exposed to single UV-AB episode at a dose of 800 mJ cm^−2^, and chlorophyll a fluorescence was analyzed 24 and 48 h after stress. Ion leakage was also used to analyze cell death 48 h after UV-AB stress as described before (Burdiak *et al.*, 2015) with the following modification: total ion leakage was evaluated after freezing at −80 °C overnight.

### Herbicide treatments

Leaf disks cut from 4-to 5-week-old plants were treated with 62.5 μM of 3-(3,4 dichlorophenyl)-1,1-dimethylurea (DCMU) and 5 μM of N,N’-dimethyl-4,4’-bipyridinium dichloride (methyl viologen, MV) with 0.05% Tween-20 and were kept in the growing chamber under ambient light between measurements. The maximum efficiency of PSII (*F*v’/*F*m’) of leaf disks was measured using FluorCam 800MF (Photon Systems Instruments, Czech Republic). As a control, we used leaf disks incubated in identical conditions without herbicides.

### RNA isolation and transcriptome profiling

For RNA isolation, only young (not fully developed) leaves were used. In the middle of the photoperiod, the leaves were detached and frozen in liquid nitrogen. For HL treatment, whole rosettes were illuminated with blue light (length 1100 μmol photons m^−2^ s^−1^) for 1 h. Immediately following the treatment, young leaves were detached and frozen in liquid nitrogen. Leaf samples represent three biological replicates, and each contained leaves from at least three individual plants. Frozen leaves were ground using mortar and pestle in liquid nitrogen, and RNA was isolated using Spectrum™ Plant Total RNA Kit (Sigma). The quality and quantity of total RNA were evaluated using Experion™ StdSens RNA Kit (Bio-Rad), and the samples with RQI value higher than 9.0 were used for RNA-seq library construction using TruSeq RNA Sample Prep Kit v2 (Illumina). RNA-seq libraries were sequenced in 100 bp paired-end reads using Illumina HiSeq 4000. RNA-seq library preparation and sequencing were conducted by Macrogen (Seoul, Korea). Reads were pseudomapped to Arabidopsis *thaliana* cDNAs (Ensembl, TAIR 10, release 35) using Salmon software (Patro *et al.*, 2017). Transcript-level abundances were imported into R using tximport (Soneson *et al.*, 2016) and analyzed using DESeq2 package (Love *et al.*, 2014). Significantly enriched GO terms were identified in up-/downregulated gene sets using PANTHER Classification System at http://geneontology.org/.

### Availability of supporting data

RNA-seq data are provided at (uploaded upon manuscript acceptance).

### Quantitative real-time PCR

RNA for qRT-PCR was isolated as described above. RNA was treated with DNase (TURBO DNA-free™ Kit, Thermo Fisher Scientific) to remove residual DNA. Next, cDNA was synthesized from 1 μg of total RNA using High-Capacity cDNA Reverse Transcription Kit (Applied Biosystems). PCRs were run on a 7500 Fast Real-Time PCR System (Applied Biosystems) using PerfeCTa™ SYBR® Green FastMix™, Low ROX™ (Quanta BioSciences) according to the manufacturer’s instructions. All primers used in qRT-PCR are listed in Table S2. The amplification efficiency of primers was calculated on the basis of the amplification curve using LinRegPCR software (Ramakers *et al.*, 2003). Relative expression was calculated using EasyqpcR package (http://www.bioconductor.org/packages/release/bioc/html/EasyqpcR.html) with *PP2AA3* and *TIP41L* as reference genes.

### Northern blot and polysome analysis

Northern blot analysis and polysome analysis were done as described previously (Fristedt *et al.*, 2014) but using Hybond-N membranes (GE Healthcare Life Sciences). Probes were amplified from total plant DNA using gene-specific primers (Table S2), radioactively labeled using the Megaprime DNA Labeling System (GE Healthcare Life Sciences), and hybridized at 65 °C.

### Thermotolerance tests

Arabidopsis seeds of WT, *cia2–2*, *cil-1*, and *cia2–2 cil-1* plants were surface sterilized by soaking in a 50% (v/v) commercial bleach solution for 7 min. The seeds were then rinsed five times with sterile water and resuspended in 1 ml of 0.1% (w/v) agarose solution. The seeds were stratified for 2 days at 4 °C. They were then placed evenly onto plates with 35 ml of ½ MS medium with agar at pH 5.8. Each plate was divided into four equal parts, one part per genotype, and approximately 25 seeds of every genotype were sown on a single plate. The plates were parafilm sealed to prevent dehydration of growing plants. The seeds were germinated and cultivated for 7 days under standard growth conditions.

After 7 days of *in vitro* cultivation, the seedlings were subjected to thermotolerance assay according to a previously described procedure (Silva-Correia *et al.*, 2014). Heat treatment was performed under ambient laboratory light after 3 h of light photoperiod. Parafilm-sealed plates with grown seedlings were set in a water bath preheated to 45 °C for 20, 30, or 40 min. After the treatment, the plates were transferred again to the growth chamber where the plants were cultivated for subsequent 7 days.

The survival rate of seedlings of individual genotypes was evaluated from the heat-treated plants’ ability to produce new green leaves 7 days after the treatment. The plants were imaged, and the proportion of seedlings survived after heat treatment and all germinated plants was calculated for each genotype at every timepoint.

### Accession numbers

The following gene names are used in Figure 5: *Rps6 (AT1G64510), Rpl11 (AT1G32990), Rpl18 (AT1G48350), Rpl28 (AT2G33450), Rpl3 (AT2G43030), Rpl21 (AT1G35680), PSRP2 (AT3G52150), Rpl9 (AT3G44890), Rps17 (AT1G79850), Rps5 (AT2G33800), Rps10 (AT3G13120), Rpl17 (AT3G54210), Rpl13 (AT1G78630), PSRP4 (AT2G38140), Rps9 (AT1G74970), Rps21 (AT3G27160), rpl2 (ATCG01310 and ATCG00830), Rps20 (AT3G15190), Rpl19.1 (AT4G17560), Rpl19.2 (AT5G47190), Rpl29 (AT5G65220), rpl32 (ATCG01020), DG1 (AT5G67570), PPR2 (AT3G06430), SVR7 (AT4G16390), PPR5 (AT4G39620), EMB2279 (AT1G30610), PDM1/SEL1 (AT4G18520) PPR596 (AT1G80270), HCF152 (AT3G09650), PSRP1 (AT5G24490), PBR1 (AT1G71720), IF3–4 (AT4G30690), IF3–2 (AT2G24060), cpHSC70–2 (AT5G49910), and SVR3 (AT5G13650).*

## Acknowledgments

We thank Prof. Ralph Bock for *psrp5-R1* and *rps17–1* seeds and Prof. Dario Leister for *prpl11–1* and *prps1–1* seeds. We thank Dr. Peter Robert Kindgren for help with *PPR* gene analysis and Dr. Anna Kozłowska-Makulska for technical help during the cHL screening. P.G., P.B., J.M., M.Z., and S.K. acknowledge financial support from the Polish National Science Center (Narodowe Centrum Nauki; OPUS6, UMO-2013/11/B/NZ3/00973 and MAESTRO6 UMO-2014/14/A/NZ1/00218 given to S.K.). L.B.S. acknowledges financial support from the Independent Research Fund Denmark (Danmarks Frie Forskningsfond; 7014–00322B). C.W. acknowledges financial support from the Academy of Finland (Decision 294580).

## Short legends for Supporting Information

**Table S1.** List of genes selected from targeted reverse genetic screen for subcellular localization experiments.

**Table S2.** List of primers used in this study.

**Figure S1.** Examples of results obtained from the initial reverse genetic screening in UV-AB and high-light experiments.

**Figure S2.** Subcellular localization of CIA2^1–100^:YFP and 100 aa N-terminal part of CIA2 fused to yellow fluorescence protein (YFP) at C termini.

**Figure S3.** Isolation and characterization of *cia2–2 cil-2* double mutant.

**Figure S4.** High-light (HL) and UV-AB susceptibility of *cia2–2 cil-2*.

**Figure S5.** CIA2 and CIL are required for optimal photosynthesis in Arabidopsis.

**Figure S6.** Analysis of rRNA and NPQ in plastid translation mutants.

**Data S1.** List of T-DNA mutants used in targeted reverse genetic screen.

**Data S2.** List of genes differentially expressed in *cia2–2 cil-1* compared to Col-0.

## Author contributions

P.G., C.W., and S.K. formulated the hypothesis and conceived the research plan. P.G., P.B., L.B.S., M.Z., and J.M. performed the experiments; P.G., P.B., and L.B.S. analyzed the data; P.G., C.W., and S.K. supervised the analysis; and P.G., P.B., C.W., L.B.S., and S.K. wrote the article, with contributions from all authors.

