## Supplementary information for "Dual-targeted transcription factors are required for optimal photosynthesis and stress responses in *Arabidopsis thaliana*"

**Table S1.** List of genes selected from targeted reverse genetic screen for subcellular localization experiments.

| **Gene** | **TF family** | **cTP**[a] | **NLS/NES**[b] | **localization**[c] |
|---|---|---|---|---|
| AT2G02080 | C2H2 | 0,888 | - | nuc |
| AT5G57180 | Orphans | 0,92 | yes | cp, nuc |
| AT3G06590 | bHLH | 0,973 | yes | - |
| AT2G34450 | HMG | 0,928 | yes | nuc |
| AT4G14700 | PHD | 0,957 | yes | nuc |
| AT4G00270 | GeBP | 0,876 | yes | nuc |
| AT5G01305 | bHLH | 0,982 | yes | - |
| AT5G43540 | C2H2 | 0,892 | yes | nuc |

[a]Chloroplast target peptide (cTP) score based on TargetP (http://www.cbs.dtu.dk/services/TargetP/). [b]presence of predicted nuclear localization/export signals (NLS/NES). [c]*in vivo* localization of proteins fused to YFP and expressed in Arabidopsis seedlings.

**Table S2.** List of primers used in this study.

| Application | Name | Sequence (5' - 3') |
|---|---|---|
| Genotyping | LBb1_3 | ATTTTGCCGATTTCGGAAC |
| | LB3 | TAGCATCTGAATTTCATAACCAATCTCGATACAC |
| | pSKTAIL-L3 | ATACGACGGATCGTAATTTGTCG |
| | DS5_3 | CGGTCGGTACGGGATTTTCC |
| | SALK_004037_LP | CTTCACCATTCGCTCACTCTC |
| | SALK_004037_RP | TTTTCGATTTCACCACCGTAG |
| | SGT49_LP | TGTAAAACCTCACTCGTTGGC |
| | SGT49_RP | CGTTAGAGCTTCGATTCCATG |
| | SALK_045340_LP | TACCTTCATTCGAGGACGTTG |
| | SALK_045340_RP | GAGGAGAAATCCGACGGTAAG |
| | SAIL_228_C01_LP | CCTCTCTGTTTTCGCATCAAC |
| | SAIL_228_C01_RP | CTAAACGAAACCTTCCGTTCC |
| | SK14786_LP | GTTTCAAAGGGGAGATTCGAC |
| | SK14786_RP | GGTAAGGAGTCACCGTTCTCC |
| Northern-blot and polysome analysis | psbDfor | TCGCTTTAGGGGGTTGGTTC |
| | psbDrev | GTGACCAAAAGCGGTTAGCG |
| | 16Sfor | TCTCATGGAGAGTTCGATCCT |
| | 16Srev | AAAGGAGGTGATCCAGCC |
| | 23Sfor | TTCAAACGAGGAAAGGCTTACG |
| | 23Srev | AGGAGAGCACTCATCTTGG |
| qPCR | PP2AA3_qPCR_F | TAACGTGGCCAAAATGATGC |
| | PP2AA3_qPCR_R | GTTCTCCACAACCGCTTGGT |
| | TIP41L_qPCR_F | GAACTGGCTGACAATGGAGTG |
| | TIP41L_qPCR_R | ATCAACTCTCAGCCAAAATCG |
| | CIA2_a_F | ATTTTGGCACCCTCCTTTTT |
| | CIA2_a_R | CAAATCTCCCCCTCATCAAA |
| | CIA2_b_F | GTGTTTAAGCAGCGGAGGAG |
| | CIA2_b_R | AGGATGATGGTGGTGGTGAT |
| | CIL_a_F | GCCGTCAAACAACACTCCTT |
| | CIL_a_R | GCCTTCTCGTGGAGATTGAG |
| | CIL_b_F | TCTCCACCGCTTACCCTAAA |
| | CIL_b_R | TGGAATCGTTGGATTGAACA |
| | RPS6_qPCR_F1 | GTGACTTGAATGAAGAAAGGATGA |
| | RPS6_qPCR_R1 | GGTAATCTTGTACTTTCTGGTTAATGC |
| | RPL11_qPCR_F1 | CCTCAACTCCGAGATTTCTCAC |
| | RPL11_qPCR_R1 | AAGCAAGTTTGATAACTCCCACA |
| | RPL18_qPCR_F1 | CCTCTGGACCAACCATTGAG |
| | RPL18_qPCR_R1 | GTGATACCTTTCTCCAAGCAAGA |
| | RPL28_qPCR_F1 | CCGTATCTCCCTTCCTTCGT |
| | RPL28_qPCR_R1 | AACTTTGTTTGCTCTGTTTGCTT |
| | RRN23S_A_F | GATGGCGAGAGTCCAGTAGC |
| | RRN23S_A_R | CAAGGTGGTCCTTGCTGATT |
| | RRN23S_B_F | GCTAAGGCCCCTAAATGACC |
| | RRN23S_B_R | GTGGCTGCTTCTAGGCAAAC |
| | RRN23S_C_F | GGCGTTAGAGCATTGAGAGG |
| | RRN23S_C_R | TGGGCACGATAACTGGTACA |
| | RRN23S_AB_F | GGGTGACCGATAGCGAAGTA |
| | RRN23S_AB_R | CTTGGGAGCTTACGGTTTCA |
| | RRN23S_BC_F | GGCAAAATAGCCCCGTAACT |
| | RRN23S_BC_R | CGGAGACCTGTGTTTTTGGT |

| | | |
|---|---|---|
| | RRN16S_F | AGCGTTATCCGGAATGATTG |
| | RRN16S_R | GATTTGACGGCGGACTTAAA |
| | HSFA2_qPCR_F | TGGGATTCTCATAAGTTCTCAACA |
| | HSFA2_qPCR_R | TGGATCAATCTTTCTGAATCCAT |
| | HSP70-4_qPCR_F | CTGACAGCGAGCGTCTCAT |
| | HSP70-4_qPCR_R | GGATCACTGTATCTTCTTCCGATT |
| USER clonning | CIA2-pLIFE-F1 | GGCTTAA/ideoxyU/ATGTCGGCGTGTTTAAGCAGC |
| | CIA2-pLIFE-R1 | GGTTTAA/ideoxyU/CCTCTTTGTCCACTTGGAGTGCTC |
| | CIA2-pLIFE-R2 | GGTTTAA/ideoxyU/CCAGAGTTTGATGAAGAGTGAGT |
| Gateway clonning | AT2G02080_LP | CACCATGTCGTCATCATCATATAAC |
| | AT2G02080_RP | ACCTCTTCCAAATGGATAATTTTGC |
| | AT5G57180_LP | CACCATGTCGGCGTGTTTAAGCA |
| | AT5G57180_RP | TCTTTGTCCACTTGGAGTGCTCTC |
| | AT3G06590_LP | CACCATGGCGTCTCTGATCTCAGAT |
| | AT3G06590_RP | AATCGGTGGAGGAGCTGAGCC |
| | AT2G34450_LP | CACCATGACGAAGAGAGCTCCCAA |
| | AT2G34450_RP | TTCAGAATAGTCTGAGTCGGTCTCTG |
| | AT1G62310_LP | CACCATGGATTCTGGAGTTAAATTG |
| | AT1G62310_RP | AAGAGATAAAAGACTTGCCTC |
| | AT4G14700_LP | CACCATGGCTTCTTCTCTGAGTTC |
| | AT4G14700_RP | CAAGTAATTGGCCAACCATGGAAG |
| | AT4G00270_LP | CACCATGGTGACTCCGAAGCAGAT |
| | AT4G00270_RP | ACTATCATTAGCTGCCTCTGCAAG |
| | AT3G50700_LP | CACCATGCCGGTAGATTTAGATAAC |
| | AT3G50700_RP | TGATTTTCTTCTACTAATGTCTTTCCC |
| | prom CIA2_LP (pDONR B4B1) | GGGGACAACTTTGTATAGAAAAGTTGCTTCTAGCTACAATTAATCAGGG |
| | prom CIA2_RP (pDONR B4B1) | GGGGACTGCTTTTTTGTACAAACTTGTTTTTTTTGTCCGGTGAAATCG |
| | CDS CIA2_LP (YFP_C) | GGGGACAAGTTTGTACAAAAAAGCAGGCTCA ATGTCGGCGTGTTTAAGCAGC |
| | CDS CIA2_RP (YFP_C) | GGGGACCACTTTGTACAAGAAAGCTGGGTTGACATCGACAGCTTCCGATCC |
| | CDS CIA2_LP (YFP_N) | GGGGACAAGTTTGTACAAAAAAGCAGGCTCA ATGTCGGCGTGTTTAAGCAGC |
| | CDS CIA2_RP (YFP_N) | GGGGACCACTTTGTACAAGAAAGCTGGGTTATTGACATCGACAGCTTCCGA |

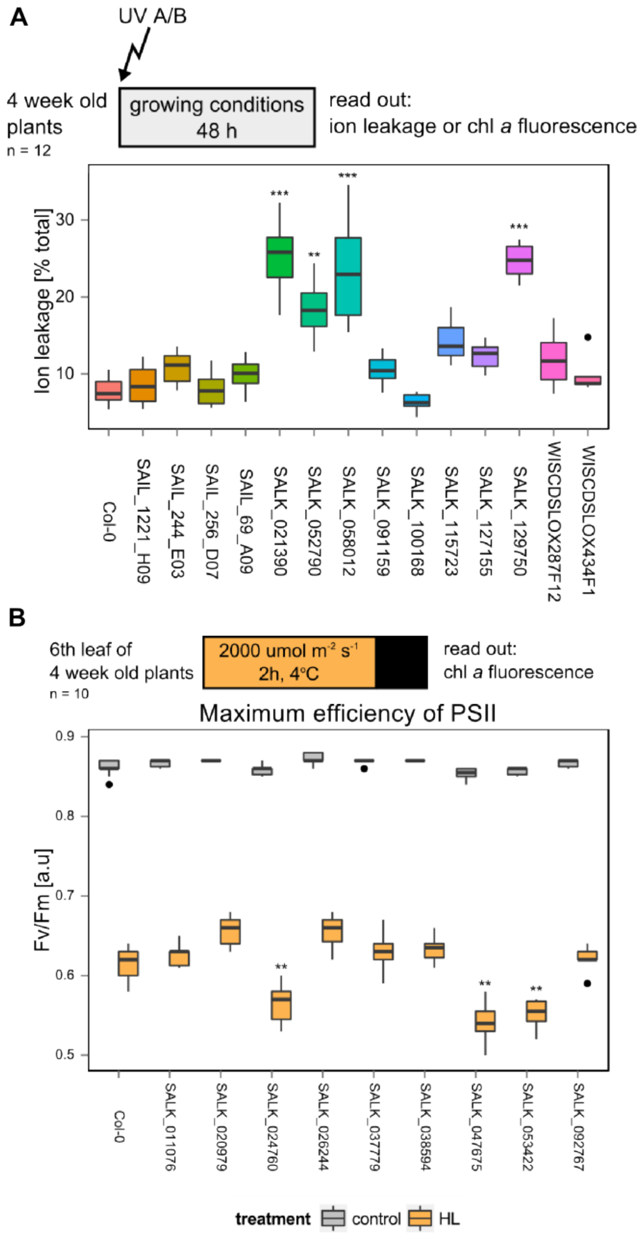


**Figure S1.** Examples of results obtained from the initial reverse genetic screening in UV-AB **(a)** and high-light (HL) **(b)** experiments. In the upper part of each panel, a graphical overview of the experimental setup is presented. Statistical significance (ANOVA and Tukey HSD test) is shown relative to Col-0 (*$p < 0.05$; **$p < 0.01$; ***$p < 0.001$).

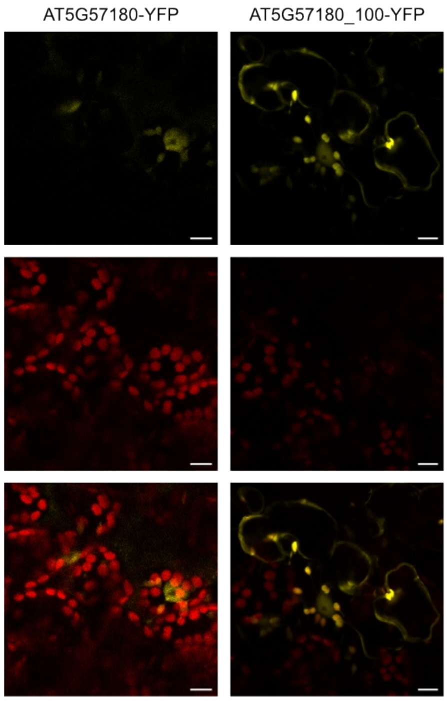


**Figure S2.** Subcellular localization of CIA2 and 100 aa N-terminal part of CIA2 fused to yellow fluorescence protein (YFP) at C termini. Fusion proteins were expressed in Arabidopsis seedlings using transient Agrobacterium transformation. Top: YFP fluorescence, middle: chlorophyll autofluorescence, and bottom: merged channels. Scale bar represents 10 µm.

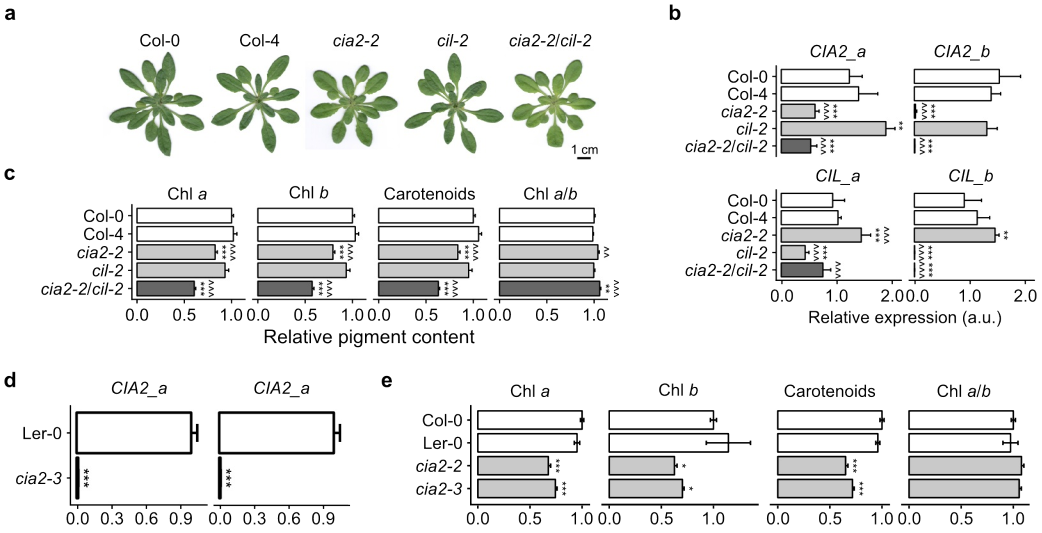


**Figure S3.** Isolation and characterization of *cia2–2 cil-2* double mutant and *cia2–3* mutant. (**a**) Phenotypes of 4-week-old plants grown in long-day conditions. *cia2–2* and *cia2–2 cil-2* plants were paler than the rest of analyzed genotypes; thus, the content of photosynthetic pigments was measured and shown in (**c**). Photosynthetic pigments were extracted from 15 plants (n = 15), bars indicate mean values ±SEM. (**b**) Expression of *CIA2* and *CIL* in analyzed genotypes was measured using qRT-PCR with gene-specific primers. Bars indicate mean values ±SD of three independent biological replicates. (**b-c**) Statistical significance (ANOVA and Tukey HSD test) is shown relative to Col-0 (*$p < 0.05$; **$p < 0.01$; ***$p < 0.001$) and Col-4 (^$p < 0.05$; ^^$p < 0.01$; ^^^$p < 0.001$). (**d**) Expression of *CIA2* in Col-0 and *cia2–3*. Bars indicate mean values ±SD of two biological replicates (each consists of at least two rosettes). (**e**) Photosynthetic pigment content in *cia2–3* mutant (Ler-0 background). Col-0 and *cia2–2* are included for comparison. Bars indicate mean values ±SEM of six biological replicates (n = 6). (**d-e**) Statistical significance (ANOVA and Tukey HSD test) is shown relative to the corresponding WT (*$p < 0.05$; **$p < 0.01$; ***$p < 0.001$).

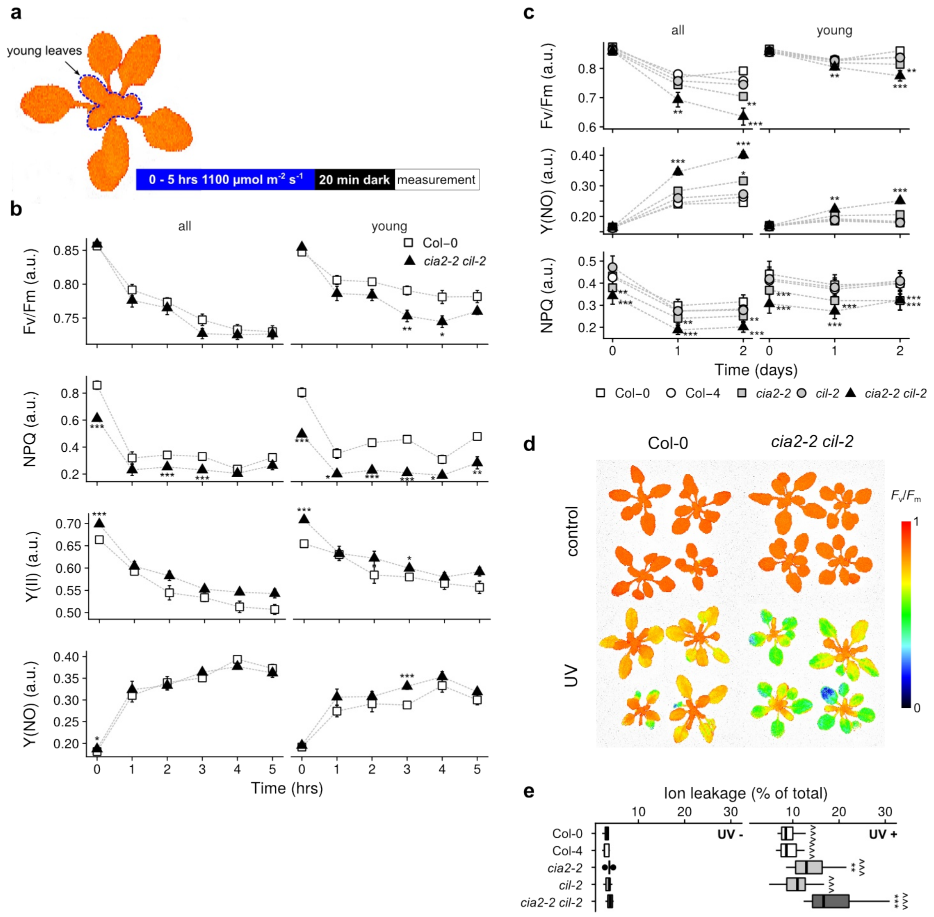


**Figure S4.** High-light (HL) and UV-AB susceptibility of *cia2–2 cil-2*. (**a**) Representation of Arabidopsis rosette with the young leaves marked with blue, dashed line. (**b**) Photosynthetic parameters of plants exposed to blue HL (1100 µmol m$^{-2}$ s$^{-1}$) for a specific period of time (n = 4–8 plants). $F_v/F_m$–maximum efficiency of PSII; NPQ–nonphotochemical quenching; Y(II)–operating efficiency of PSII; Y(NO)–nonregulated energy dissipation. (**c**) Photosynthetic parameters of plants exposed to UV-AB. Each point represents mean ±SEM of at least four plants. (**d**) $F_v/F_m$ in Col-0 and *cia2–2 cil-2* exposed to UV-AB. (**e**) Ion leakage of control (UV-, n = 5) and UV-AB-treated (UV +, n ≥ 17) plants is shown as a percentage of total ion leakage. Statistical significance (ANOVA and Tukey HSD test) is shown relative to Col-0 (**p* < 0.05; ***p* < 0.01; ****p* < 0.001) and to control conditions (^^^*p* < 0.001).

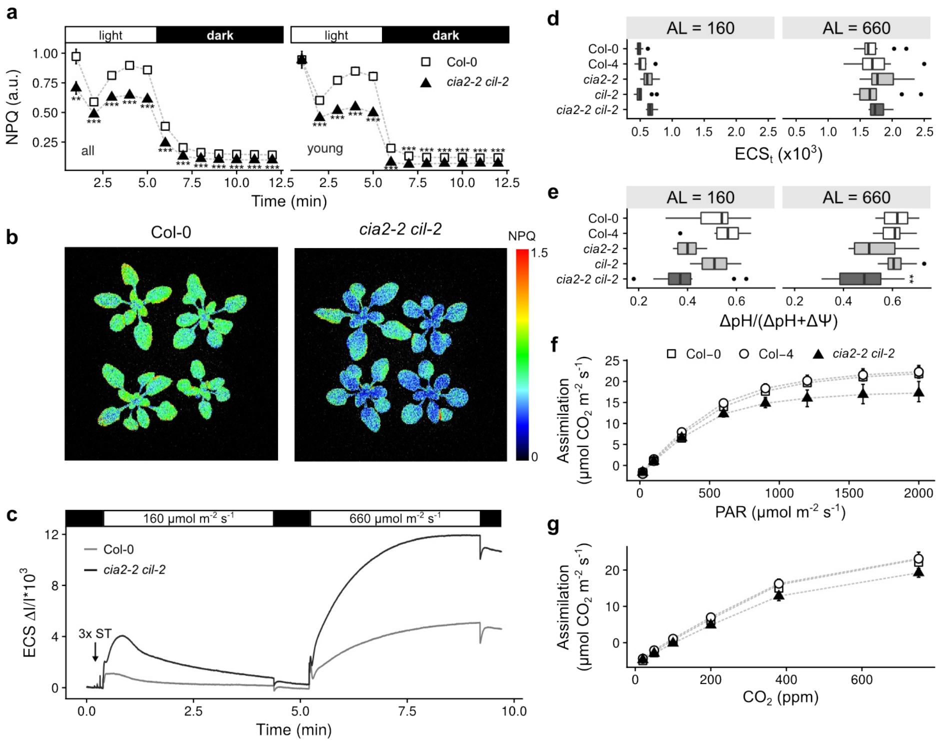


**Figure S5.** CIA2 and CIL are required for optimal photosynthesis in Arabidopsis. (**a**) Nonphotochemical quenching (NPQ) in analyzed genotypes. Points represent mean ±SEM (n = 4) of the whole rosette and only young leaves. (**b**) NPQ of Col-0 and *cia2–2 cil-2*. (**c**) Analysis of electrochromic pigment shift (ECS, P515) at 160 and 660 µmol $m^{-2}$ $s^{-1.}$ of actinic light. For simplicity, only Col-0 and *cia2–2 cil-2* are shown. ST–single turnover flash. Total ECS ($ECS_t$) (**d**) and ΔpH (**e**) in analyzed genotypes at 160 and 660 µmol $m^{-2}$ $s^{-1}$. Box plots represent values of 10 independent plants. (**f**) $CO_2$ assimilation as a function of light intensity. (**g**) $CO_2$ assimilation as a function of $CO_2$ concentration. In (**f**) and (**g**), values represent mean ±SEM of 7–9 plants. Statistical significance (ANOVA and Tukey HSD test) is shown relative to Col-0 (*$p$ < 0.05; **$p$ < 0.01; ***$p$ < 0.001).

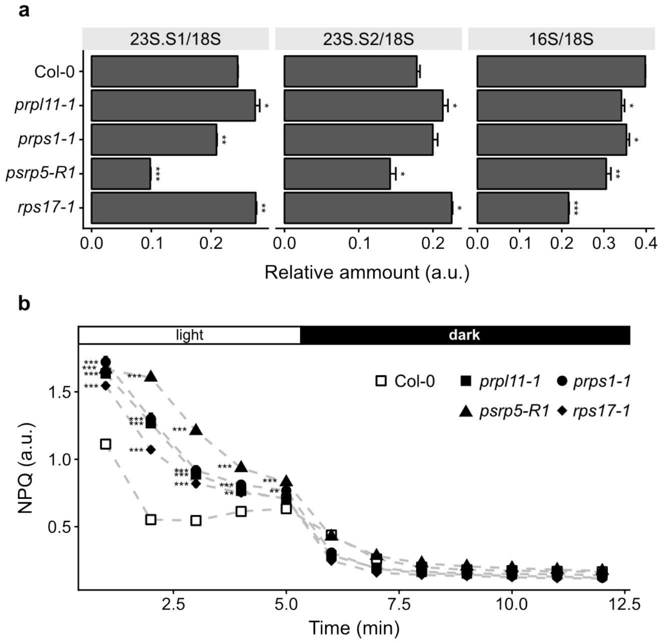


**Figure S6.** Analysis of rRNA and NPQ in plastid translation mutants. Both experiments were performed on 19-day-old plants grown in standard conditions. (**a**) Relative abundance of precursor 23S and 16S rRNAs was measured using capillary electrophoresis, and mean values ±SEM are shown (n = 2, each consists of at least three plants). (**b**) NPQ induction curve (150 µmol photons $m^{-2}$ $s^{-1}$). Plants were illuminated with actinic light for 5 min followed by recovery for 7 min. Points represent mean values ±SEM (n ≥6). Statistical significance (ANOVA and Tukey HSD test) is shown relative to Col-0 (*$p < 0.05$; **$p < 0.01$; ***$p < 0.001$).
